# Cortical and subcortical mapping of the allostatic-interoceptive system in the human brain using 7 Tesla fMRI

**DOI:** 10.1101/2023.07.20.548178

**Authors:** Jiahe Zhang, Danlei Chen, Philip Deming, Tara Srirangarajan, Jordan Theriault, Philip A. Kragel, Ludger Hartley, Kent M. Lee, Kieran McVeigh, Tor D. Wager, Lawrence L. Wald, Ajay B. Satpute, Karen S. Quigley, Susan Whitfield-Gabrieli, Lisa Feldman Barrett, Marta Bianciardi

## Abstract

The brain continuously anticipates the energetic needs of the body and prepares to meet those needs before they arise, called allostasis. In support of allostasis, the brain continually models the sensory state of the body, called interoception. We replicated and extended a large-scale system supporting allostasis and interoception in the human brain using ultra-high precision 7 Tesla functional magnetic resonance imaging (fMRI) (*N =* 90), improving the precision of subgenual and pregenual anterior cingulate topography combined with extensive brainstem nuclei mapping. We observed over 90% of the anatomical connections published in tract-tracing studies in non-human animals. The system also included regions of dense intrinsic connectivity broadly throughout the system, some of which were identified previously as part of the backbone of neural communication across the brain. These results strengthen previous evidence for a whole-brain system supporting the modeling and regulation of the internal milieu of the body.

## Introduction

A brain efficiently regulates and coordinates the systems of the body as it continually interfaces with an ever-changing and only partly predictable world. Various lines of research, including tract-tracing studies of non-human animals (e.g., 1, 2), discussions of predictive processing (3–6), and research on the central control of autonomic nervous system function (7–11), all suggest the existence of a unified, distributed brain system that anticipates the metabolic needs of the body and prepares to meet those needs before they arise, a process called allostasis (10; for recent reviews, see 11, 12). Allostasis is not a condition or state of the body — it is the process by which the brain efficiently coordinates and regulates the various systems of the body (12). Just as somatosensory and other exteroceptive sensory signals are processed in the service of skeletomotor control, the brain is thought to model the internal sensory conditions of the body (i.e., the internal milieu) in the service of allostasis, a process known as interoception (15–18).

Using resting state functional magnetic resonance imaging (fMRI) in three samples totalling almost 700 human subjects scanned at 3 Tesla (19), we previously identified a distributed allostatic-interoceptive system consisting of two well-known intrinsic networks, the default mode network and salience networks, overlapping in many key cortical visceromotor allostatic regions that also serve as ‘rich club’ hubs that have been implicated as the “backbone” for neural communication throughout the brain (**Figure 1A**). Our investigation was guided by the anatomical tracts identified in published studies of macaques and other non-human mammals (see Table 2 in (19)). This first study was more cortically focused, examining the functional connectivity of primary interoceptive cortex spanning the dorsal mid and dorsal posterior insula (dmIns/dpIns), as well as key allostatic regions in the cerebral cortex that are directly connected to the brainstem regions that are known to be responsible for controlling the motor changes in the viscera (i.e., visceromotor cortical regions), such as the anterior midcingulate cortex (aMCC), pregenual anterior cingulate cortex (pACC), subgenual anterior cingulate cortex (sgACC), and agranular insular cortex (also known as ventral anterior insula, or vaIns, which is also posterior orbitofrontal cortex), as well as the dorsal sector of the amygdala (dAmy) containing the intercalated bodies and the central nucleus (**Figure 1A**). Our 3T analysis yielded a replicable, integrated system consisting of two well-known intrinsic networks, in addition to primary interoceptive cortex. We did explore some aspects of the system’s subcortical extent, including the thalamus, hypothalamus, hippocampus, ventral striatum, periaqueductal gray (PAG), parabrachial nucleus (PBN) and nucleus tractus solitarius (NTS), all regions known to play a role in control of the autonomic nervous system, the immune system, and the endocrine system (e.g., 18–24), but our ability to more extensively map the midbrain and brainstem extents of the system were limited by our use of 3T imaging.

**Figure 1.**
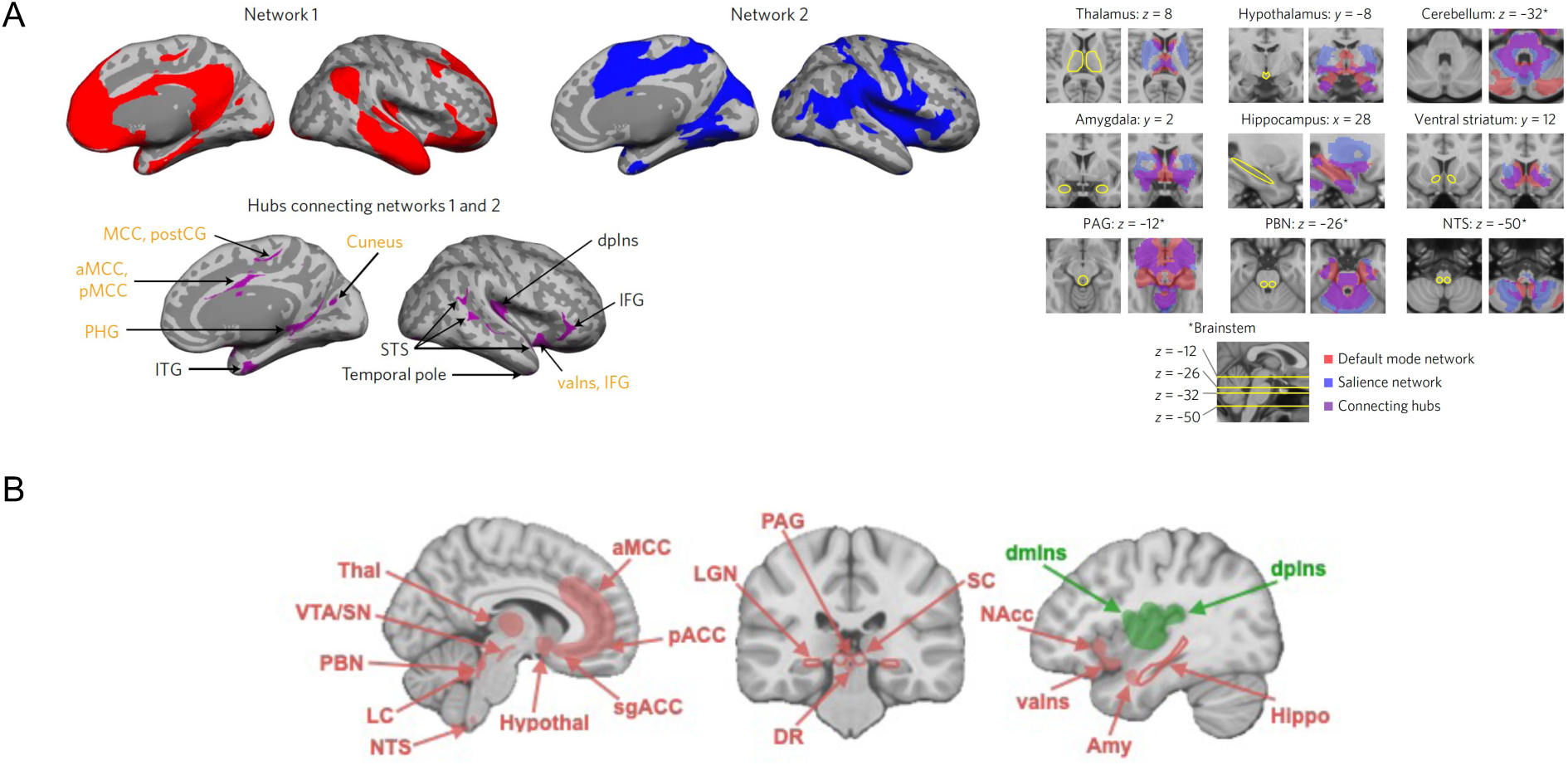
Key cortical and subcortical regions involved in interoception and allostasis. **(A)** Using 3 Tesla fMRI resting state connectivity, we showed a unified system consisting of the default mode network (in red) and salience network (in blue), which overlapped in many key cortical visceromotor allostatic regions (in purple) that also serve as ‘rich club’ hubs, in addition to a portion of primary interoceptive cortex (dpIns) (left panel) (19). We reported the system’s connectivity to some subcortical regions known to play a role in control of the autonomic nervous system, the immune system, and the endocrine system such as the thalamus, hypothalamus, hippocampus, ventral striatum, PAG, PBN and NTS (e.g., 20–26) (right panel) (19). Figures are reproduced with permission from (19). (B) Expanded set of seed regions used in the present analysis. Abbreviations: aMCC: anterior midcingulate cortex; Amy: amygdala; dmIns: dorsal mid insula; dpIns: dorsal posterior insula; DR: dorsal raphe; Hippo: hippocampus; Hypothal: hypothalamus; LC: locus coeruleus; LGN: lateral geniculate nucleus; NAcc: nucleus accumbens; NTS: nucleus of the solitary tract; pACC: pregenual anterior cingulate cortex; PAG: periaqueductal gray; PBN: parabrachial nucleus; SC: superior colliculus; sgACC: subgenual anterior cingulate cortex; SN: substantia nigra; Thal: thalamus; vaIns: ventral anterior insula; VTA: ventral tegmental area.

In the present study, we replicated and extended evidence for the allostatic-interoceptive system (**Figure 1B**) using ultra-high field (7 Tesla) MRI, which allows data acquisition with higher spatial resolution (1.1 mm isotropic), better signal-to-noise-ratio (SNR; (27–29)), and increased sensitivity in mapping functional connectivity of brainstem nuclei involved in arousal, motor and other vital processes (e.g., autonomic, nociceptive, sensory; (30, 31)). This is a particularly important effort given the increasing importance of the allostatic-interoceptive system as a tool for investigating interoception and allostasis in basic brain function both in neurotypical samples and in specific populations (e.g., (32–35)). In addition, research indicates that regions in this system are also important for a wide range of psychological domains, including cognition, emotion, pain, decision-making and perception (see Figure 5 in (19); also see (36–38)), suggesting the hypothesis that allostatic and interoceptive signals may play a more fundamental role in shaping basic brain dynamics (for discussion see (5, 39–41)).

We tested within-system functional connectivity in 90 human participants (age range = 18-40 years, mean = 26.9 years, s.d. = 6.2 years; 40 females) using a fast low-angle excitation echo-planar technique (FLEET) sequence shown to reduce artifacts and improve temporal SNR (24, 42). This approach allowed a more precise mapping of connectivity for regions with known signal issues at 3 Tesla, such as the sgACC (low SNR), amygdala (noise from adjacent veins; (43)), columns within the PAG (noise from adjacent aqueduct), and other small structures that could be particularly influenced by partial volume effects. We took advantage of recently developed, much improved and validated in-vivo brainstem and diencephalic nuclei atlases (44–48) to guide our analysis. This was crucial because our hypotheses were specifically derived from published tract-tracing studies of macaques and other non-human mammals that establish structural pathways carrying ascending interoceptive signals from the periphery, for example via the vagus nerve, to subcortical and cortical regions of the allostatic-interoceptive system (**Supplementary Table 1**). Extending (19), we more extensively examined the intrinsic connectivity of subcortical nuclei such as mediodorsal thalamus (mdThal), hypothalamus, dorsal amygdala, hippocampus, ventral striatum, PAG, PBN and the NTS (in the medullary viscero-sensory-motor nuclei complex; VSM), in addition to considering the connectivity of dorsal raphe (DR), substantia nigra (SN), ventral tegmental area (VTA), locus coeruleus (LC), superior colliculus (SC), and lateral geniculate nucleus (LGN). The DR, SN, VTA and LC are midbrain and pontine monoamine-producing nuclei that contribute to relaying the body’s metabolic status to the cortex (49). The SC and LGN are not traditionally considered to be directly involved in interoception and allostasis, but they share anatomical connections with key visceromotor regulation regions in the system (see **Supplementary Table 1**; (50–55)). For example, neurons in the intermediate and deep layers of the SC are connected to aMCC (56), hypothalamus (57, 58) and PAG (59), and have been directly implicated in skeletomotor (60, 61) and visceromotor (62, 63) actions that facilitate approach or avoidance behaviors. The SC is also thought to be a major point of sensory-motor integration and is associated with affective feelings (64, 65). The LGN receives interoceptive inputs from the PAG (52) and PBN (55, 66, 67), and shares monosynaptic connections with the hypothalamus (68) and pACC (69). We also examined connectivity patterns for subregions of the PAG, hippocampus and SC rather than as a single ROI as in (19) given their functional heterogeneity (70, 71) and differential involvement in allostasis (e.g., (72–74)).

## Results

We used a bootstrapping strategy to identify weak but reliable signals that are important when examining cortico-subcortical connections in brain-wide analyses. For each of 1000 iterations, we randomly resampled 80% of the subjects (*N* = 72) and identified, for each seed region, BOLD correlations for all voxels in the brain that survived a voxel-wise threshold of *p* < .05. We calculated discovery maps for each seed region that included both cortical and subcortical connections. We calculated the similarities in the spatial topography among all the maps and subjected each resulting similarity matrix to *k-*means clustering analysis to characterize the allostatic-interoceptive network. We expected stronger connectivity among cortical seeds compared to among subcortical seeds due to the latter’s noisier time courses and potential partial volume effects, which would result in lower correlations for smaller regions.

### Cortico-cortical intrinsic connectivity

We first examined the hypothesized functional connectivity according to published anatomical connections. As expected, we successfully replicated all of the cortico-cortical connections we previously observed with 3 Tesla imaging (**Figure 2, Supplementary Table 1**) (19). We additionally observed reciprocal intrinsic connectivity (i.e., connectivity map of one region includes a cluster in the other region and vice versa) between the lvAIns and pACC, between the sgACC and aMCC, and between the dmIns and portions of cingulate cortex (sgACC, pACC) (**Figure 2A**; see **Supplementary Figure 1** for *t-*value map based on full group), extending the allostatic-interoceptive system to include more of the anatomical connections documented in tract-tracing studies in non-human mammals (75–78). All of these observations were confirmed by seed-to-seed connectivity strength calculation (**Figure 2B**). Using evidence from the cortical maps and the seed-to-seed connectivity matrices, we confirmed 100% of the monosynaptic cortico-cortical connections documented in published tract-tracing studies in non-human animals.

**Figure 2.**
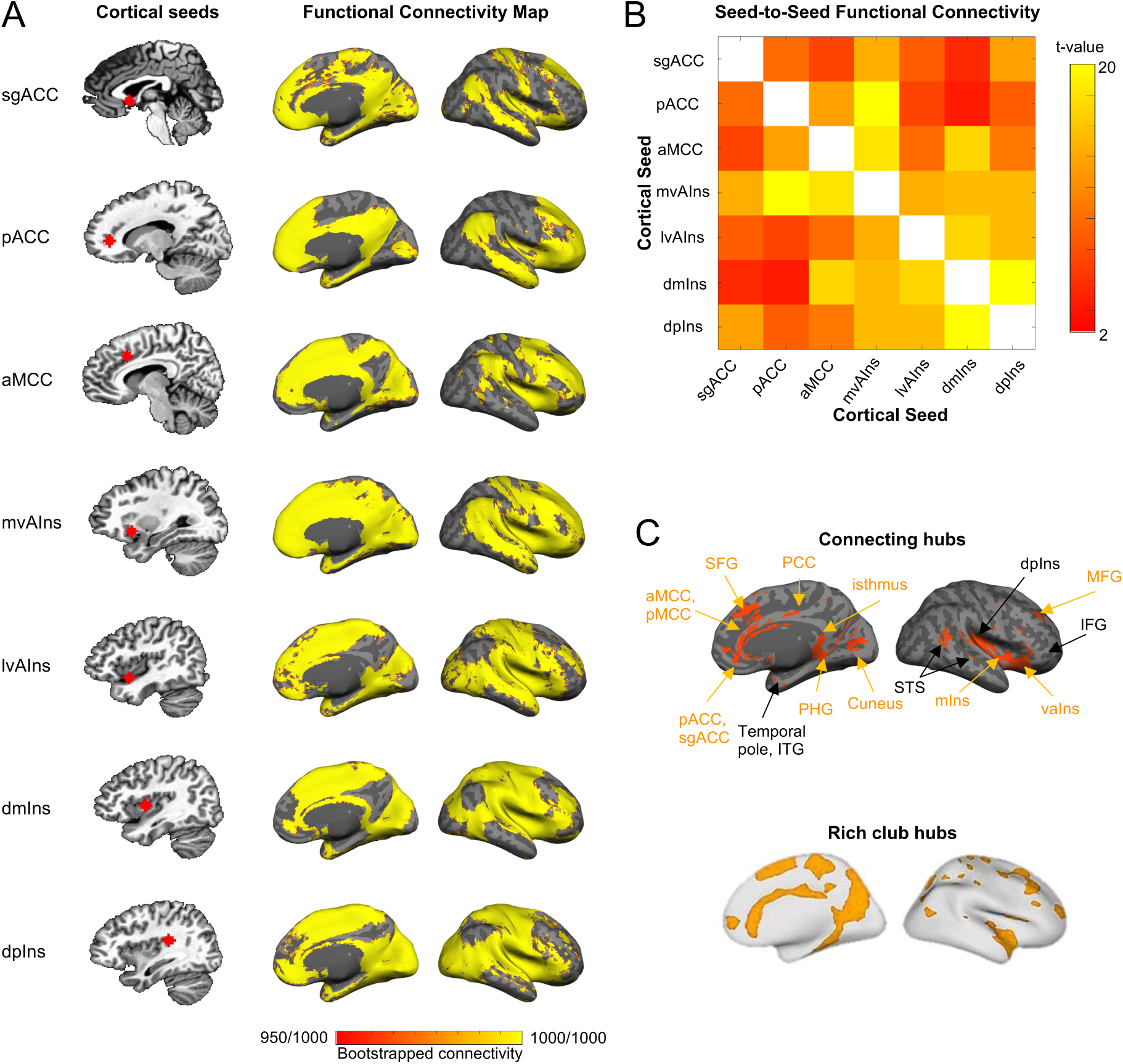
Cortico-cortical functional connectivity within the allostatic-interoceptive system. **(A)** Left column shows cortical seed locations and right column shows bootstrapped functional connectivity maps depicting all voxels whose time course was correlated (*p* < .05) with that of the seed in more than 950 iterations (out of 1000) by resampling 80% of the sample in each iteration (*N* = 72). **(B)** Seed-to-seed functional connectivity matrix shows connectivity strength between each pair of the cortical seeds (*p* < .05, uncorrected; white color indicates correlation =1; *N* = 90). **(C)** The allostatic-interoceptive system showed connecting regions in all the *a priori* interoceptive and visceromotor control regions. Connecting regions belonging to the ‘rich club’ are labeled in yellow. ‘Rich club’ hubs figure adapted with permission from (105). To avoid Type II errors, which are enhanced with the use of stringent statistical thresholds (175), we opted to separate signal from random noise using replication, according to the mathematics of classical measurement theory (176). Abbreviations: aMCC: anterior mid cingulate cortex; dpIns: dorsal posterior insula; IFG: inferior frontal gyrus; MFG: middle frontal gyrus; mIns: mid insula; pACC: pregenual anterior cingulate cortex; PHG: parahippocampal gyrus; pMCC: posterior mid cingulate cortex; PCC: posterior cingulate cortex; sgACC: subgenual anterior cingulate cortex; STS: superior temporal sulcus; vaIns: ventral anterior insula.

Next, we binarized the cortical connectivity maps for all cortical seeds (*p* < 0.05) and computed their conjunction to identify connecting cortical regions (**Figure 2C**). A *k-*means clustering analysis (optimal *k* = 2 based on the Calinski-Harabasz Criterion (79)) on the cortical maps replicated (19), such that the system included two subsystems, one corresponding to the default mode network (i.e., the dorsomedial prefrontal cortex, posterior cingulate cortex (PCC), and dorsolateral prefrontal cortex) and the other corresponding to the salience (i.e., anterior to MCC, anterior insula, supramarginal gyrus, supplementary motor area) and somatomotor networks (i.e., M1, S1, superior temporal gyrus; see details in **Supplementary Figure 2**). This procedure also allowed us to discover any regions that might be reliably included in the intrinsic connectivity of the system. We replicated all the connecting ‘hub’ regions reported at 3 Tesla in (19) (i.e., portions of anterior/posterior midcingulate cortex, inferior frontal gyrus, ventral anterior insula, dorsal posterior insula, temporal pole, inferior temporal gyrus, superior temporal sulcus, parahippocampal gyrus, and cuneus) with the exception of medial postcentral gyrus. We also newly identified as allostatic-interoceptive system ‘hubs’ the entire anterior cingulate cortex (including subgenual and pregenual extents), PCC, a greater extent of the insula (including mid insula; mIns), as well as some portions of medial superior frontal gyrus (SFG) and middle frontal gyrus (MFG). A majority of the allostatic-interoceptive system’s connecting hubs have been identified as members of the ‘rich club’ in the connectomics literature, defined as high-degree nodes showing denser interconnections among themselves than are lower degree nodes (80). The rich club hubs play a key role in global information integration across the brain and therefore may serve as the backbone for global communication in the brain (81), suggesting that allostatic and interoceptive processes may be at the core of the brain’s computational architecture.

### Subcortico-cortical intrinsic connectivity

In a new analysis enabled by newly delineated subcortical seeds (45, 48, 82) that was not possible at 3 Tesla (19), we assessed subcortico-cortical connectivity by visually inspecting cortical discovery maps of the subcortical seeds to confirm topography (**Figure 3A**; see **Supplementary Figure 3** for *t-*value map based on full group) and calculating seed-to-seed connectivity to quantify strength of connection (**Figure 3B**). Combining evidence from the cortical maps and seed-to-seed connectivity matrix, we confirmed 94% of the monosynaptic subcortico-cortical connections predicted from non-human tract-tracing studies (**Supplementary Table 1**). There were three exceptions: we did not observe significant, positive functional connectivity between PAG and dmIns/dpIns, hypothalamus and dmIns/dpIns, or PBN and sgACC, despite known anatomical connections (see **Supplementary Table 1**; (83, 84)). In some instances, averaged time courses between seeds did not correlate significantly (i.e., *gray* squares in **Figure 3B**, e.g., DR-sgACC), but connectivity clusters could nonetheless be observed in the maps (e.g., sgACC cluster in DR-seeded map). Such discrepancies can result from noisy signals within an ROI or specific sub-portions of an ROI showing significant connectivity. We tested specificity of the allostatic-interoceptive network using a region of superior parietal lobule not known for visceromotor function (19). This region only showed consistent functional connectivity to the SC (85), VSM, the hippocampus and the amygdala (**Supplementary Table 2**).

**Figure 3.**
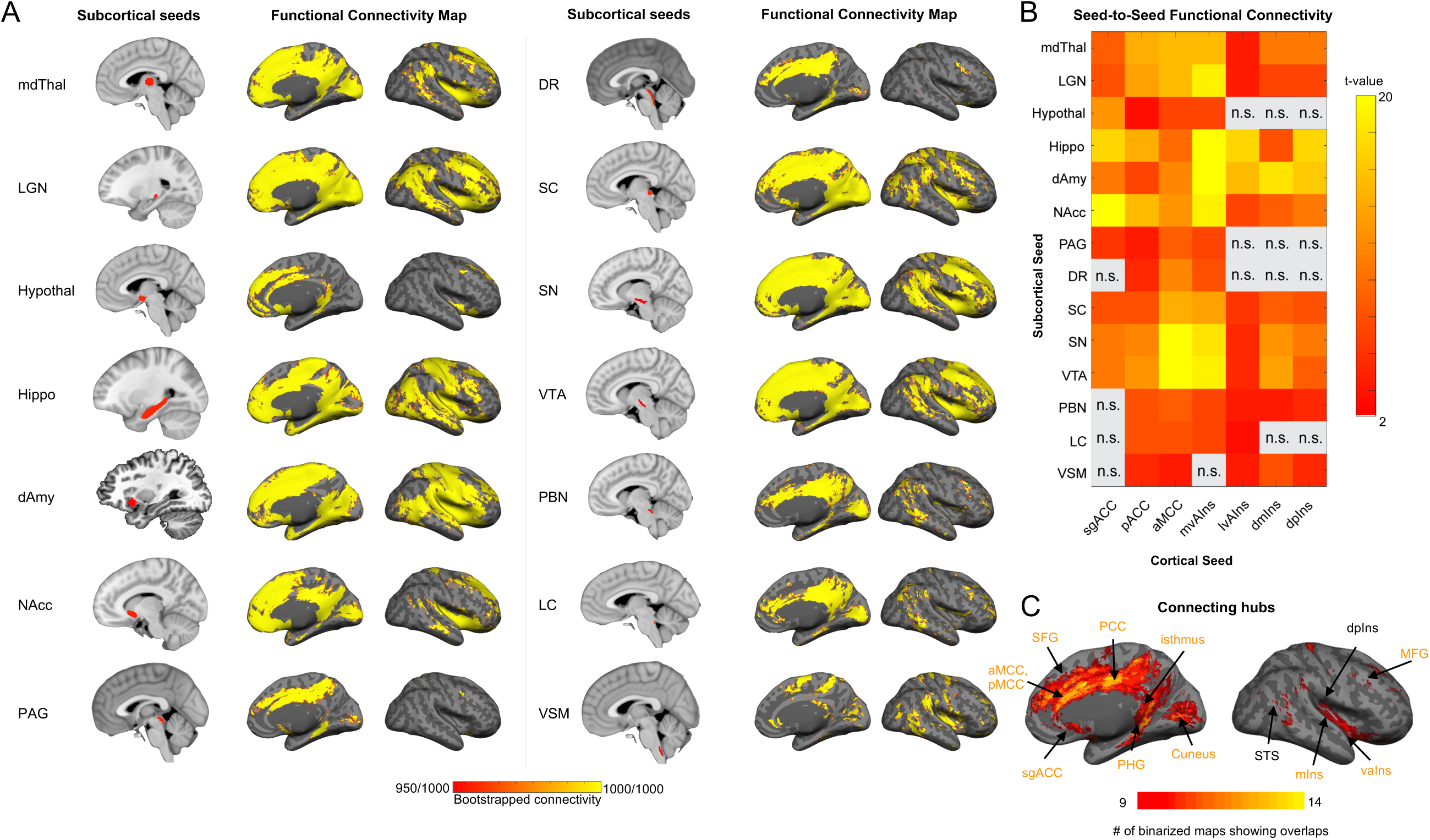
Subcortico-cortical intrinsic connectivity within the allostatic-interoceptive system. **(A)** Left column shows subcortical seed locations and right column shows bootstrapped functional connectivity discovery maps depicting all cortical voxels whose time course was correlated (*p* < .05) with that of the seed in more than 950 iterations (out of 1000) by resampling 80% of the sample in each iteration (*N* = 72). **(B)** Seed-to-seed functional connectivity matrix shows connectivity strength between pairs of subcortical and cortical seeds (*p* < .05, uncorrected; gray color indicates subthreshold correlations; *N* = 90). **(C)** Conjunction map shows the number of binarized maps (*p* < .05) with shared connecting regions (ranging from 9 to 14). Abbreviations: dAmy: dorsal amygdala; mdThal: mediodorsal thalamus; LGN: lateral geniculate nucleus; Hypothal: hypothalamus; Hippo: hippocampus; NAcc: nucleus accumbens; PAG: periaqueductal gray; DR: dorsal raphe; SC: superior colliculus; SN: substantia nigra; VTA: ventral tegmental area; PBN: parabrachial nucleus; LC: locus coeruleus; VSM: medullary viscero-sensory-motor nuclei complex corresponding to the nucleus tractus solitarius.

As with the cortico-cortical analyses, we conjoined the binarized discovery subcortico-cortical maps (*p* < 0.05) to identify the overlapping cortical connectivity between subcortical seeds (**Figure 3C**). Subcortically seeded maps showed connecting regions in hypothesized cingulate and insular regions as well as some parts of the MFG and cuneus. We examined a range of *k* values that showed similarly optimal Calinski-Harabasz Criterion (*k* = 2 to 9) (see **Supplementary Methods**). We retained *k* = 3 for its interpretability. All three clusters included cortical nodes from the default mode and salience networks. Cluster 1 included discovery maps from seeds in the lower brainstem (LC, PBN, VSM), and primarily showed connectivity to the posterior cingulate cortex, supramarginal gyrus and some medial and lateral occipital regions (**Supplementary Figure 4**). Cluster 2 included discovery maps from seeds in the upper brainstem (PAG, DR) and the hypothalamus, and showed connectivity to the aMCC and parahippocampal gyrus. Cluster 3 included discovery maps from larger seeds in the mdThal, LGN, hippocampus, dAmy, NAcc, SC, SN and VTA, and showed widespread connectivity to the dorsomedial prefrontal cortex, cingulate cortices (sgACC, pgACC, aMCC, isthmus), supplementary motor area, cuneus, insula (anterior, mid and posterior), superior frontal gyrus, central sulcus, and angular gyrus.

### Subcortico-subcortical intrinsic connectivity

With our newly delineated subcortical seeds (45, 48, 82), we also assessed subcortico-subcortical connectivity by visually inspecting subcortical maps of the subcortical seeds to confirm topography (**Figure 4A**) and by calculating functional connectivity between all subcortical seeds to quantify strength of connection (**Figure 4B**). Again, this analysis was not possible with 3 Tesla scanning as in (19). We confirmed 96% of the monosynaptic subcortico-subcortical connections that were identified in published, tract-tracing studies on non-human animals (**Supplementary Table 1**). There were three exceptions: we did not observe significant, positive functional connectivity between hypothalamus and PBN, hypothalamus and LC, or hypothalamus and VSM (including NTS), despite known anatomical connections (see **Supplementary Table 1**). In one case, averaged timecourses between the VSM and NAcc seeds did not correlate significantly (**Figure 4B**; see *gray* square in matrix), but bilateral NAcc clusters could nonetheless be observed in the VSM-seeded map. As in the subcortico-cortical maps, such discrepancies can result from noisy signals within an ROI or specific sub-portions of an ROI showing significant connectivity. Seed-to-seed connectivity strength between PAG subregions and other subcortical ROIs is displayed in **Supplementary Figure 5**. Seed-to-seed connectivity strength between hippocampal subregions and other subcortical ROIs is displayed in **Supplementary Figure 6**. Seed-to-seed connectivity strength between layers of the SC and other subcortical ROIs is displayed in **Supplementary Figure 7**. Conjoined binarized subcortical discovery maps (*p* < 0.05) indicated that all but four subcortical seeds showed overlapping connectivity: connecting regions were identified in the mdThal, LGN, hippocampus, dAmy, NAcc, PAG, DR, SC, SN and VTA but hypothalamus, PBN, LC and VSM showed less widespread and dense connectivity throughout subcortical seeds (**Supplementary Table 3**). K-means clustering analysis (*k* = 3) on the subcortical discovery maps from subcortical seeds yielded an almost identical solution as their cortical connectivity maps.

**Figure 4.**
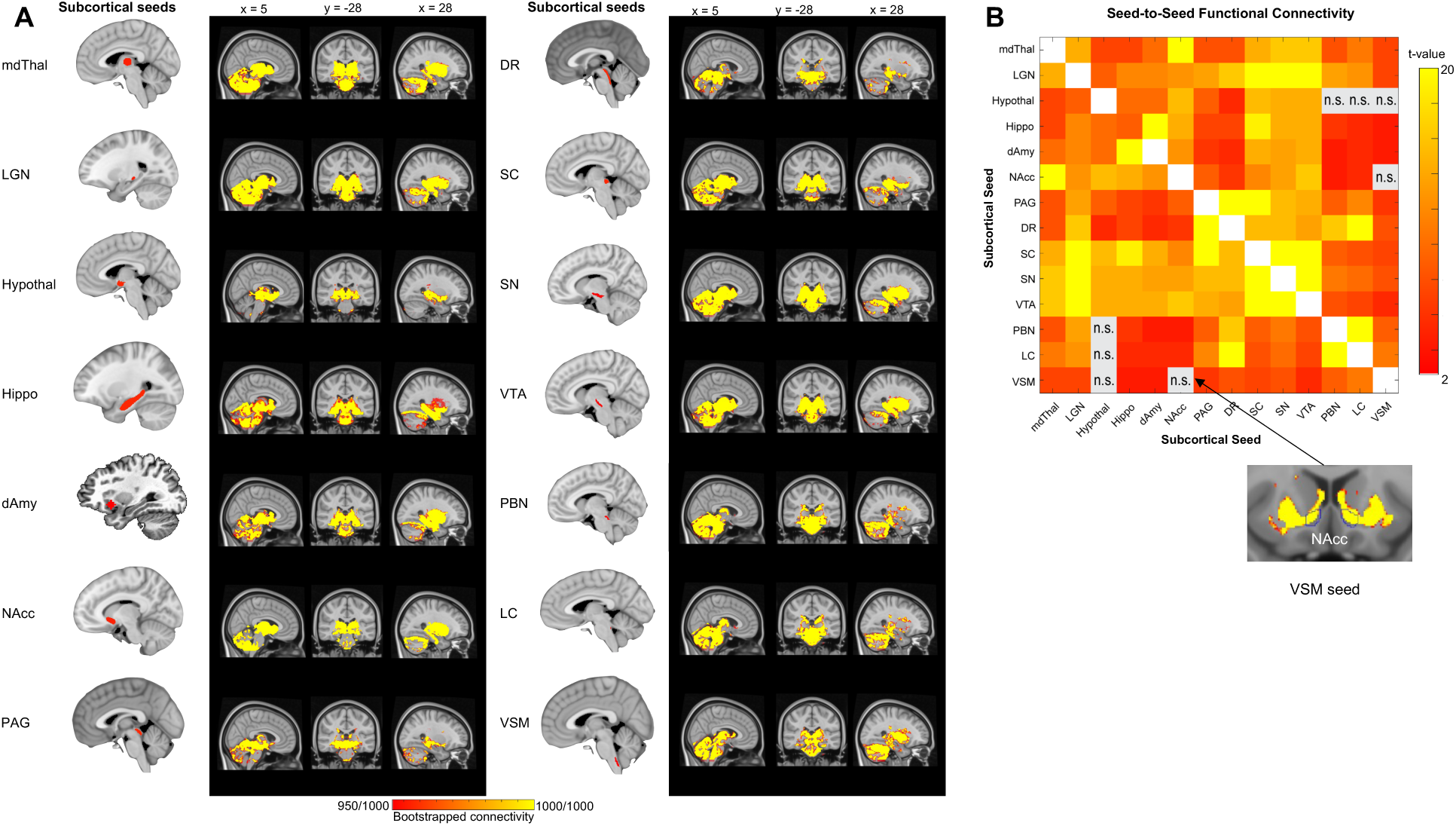
Subcortico-subcortical intrinsic connectivity within the allostatic-interoceptive system. **(A)** Left column shows subcortical seed locations and right column shows bootstrapped functional connectivity discovery maps depicting all subcortical voxels whose time course was correlated (*p* < .05) with that of the seed in more than 950 iterations (out of 1000) by resampling 80% of the sample in each iteration (*N* = 72). **(B)** Seed-to-seed functional connectivity matrix showed connectivity strength between each pair of the subcortical seeds (*p* < .05, uncorrected; white color indicates correlation =1 and gray color indicates subthreshold correlations; *N* = 90). Several seeds had functional connectivity with a subset of voxels within target ROIs, as shown by binarized maps at *p* < .05 (target ROI outline is shown in blue).

### The allostatic-interoceptive system

We observed dense interconnectivity between all the cortical and subcortical seeds included in our analysis (**Figure 5A**). Conjoined binarized discovery maps (*p* < 0.05) across both cortical and subcortical extents converged in the hypothesized allostatic-interoceptive system (**Figure 5B**).

**Figure 5.**
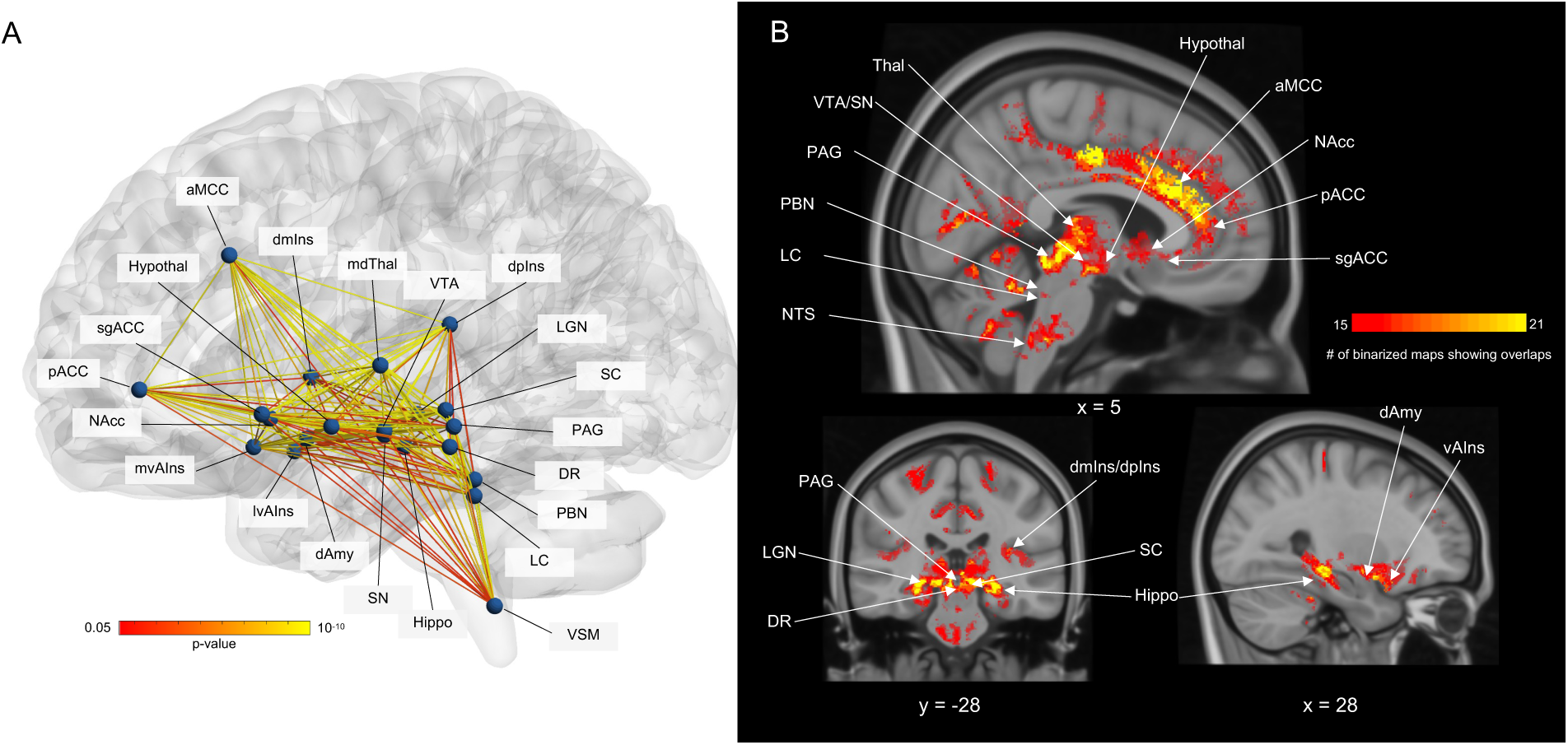
Summary of the allostatic-interoceptive system based on 7 Tesla fMRI functional connectivity. **(A)** Circuit diagram indicates dense within-system connectivity between the 21 cortical and subcortical seeds. All seeds are shown as spherical nodes located at their respective centers of gravity. Pairwise connectivity strengths between ROIs are shown as edges between nodes (ranging from *p* < .05 in red to *p* < 10^-10^ in yellow, uncorrected; *N* = 90). Nodes and edges in the glass brain were visualized using BrainNet Viewer (177) **(B)** Conjunction map shows the number of binarized maps (*p* < .05) that shared overlapping regions (ranging from 15 to 21, total number of cortical and subcortical seeds = 21).

## Discussion

Ultra-high field 7 Tesla fMRI with 1.1-mm isotropic voxel resolution combined with newly delineated 7 Tesla brainstem and diencephalic parcellations (44–48) revealed both cortical and subcortical components of an integrated allostatic-interoceptive system in humans consisting of two overlapping subsystems. Our original study using 3 Tesla fMRI (19) used 10-minute resting state scans in two subsamples of 270-280 participants each, plus a third sample of N = 41, whereas the present study involved a greater duration of resting state scan time (30 minutes total) in a sample of 90 participants. Using functional connectivity among seven cortical ROIs and 14 subcortical ROIs in humans, we verified over 90% of the anatomical connections identified in published tract-tracing studies of macaques and other non-human mammals. Our current 7 Tesla findings revealed reciprocal connectivity between sgACC/pACC and dmIns/dpIns regions previously unreported in 3 Tesla functional connectivity studies of the ACC (86–90) and the insula (91–94). The improvement in sgACC connectivity, in particular, was expected at 7 Tesla, as this region is part of the medial/orbital surface that is typically susceptible to low SNR, partial volume effects and physiological aliasing. In the current study, these effects were mitigated by higher resolution image acquisition at 7 Tesla, minimal smoothing, and more precise nuisance regression using signals from individual ventricles. We also expanded observations of the subcortical extents of the system. Several subcortical nodes (i.e., mdThal, LGN, hippocampus, dAmy, NAcc, SC, SN and VTA) showed robust connectivity with all cortical nodes whereas the smaller brainstem nuclei (i.e., PAG, DR, PBN, LC and VSM (including the NTS)) showed weaker but reliable connectivity to these nodes, consistent with other studies that examined a subset of the nodes as seeds at 3 Tesla (e.g., (49)) and 7 Tesla (e.g., (30, 95, 96)). We also observed reliable connectivity between regions that have not yet been documented as having monosynaptic connections in previous tract-tracing studies. For example, the LGN has virtually no monosynaptic connectivity with cortical nodes of the allostatic-interoceptive system according to tract-tracing studies (except for modest projections to the pACC (97)), yet we observed reliable functional connectivity between the LGN and the aMCC, mvAIns, and pACC. The LGN receives interoceptive input (67) and there is some evidence that interoceptive signals gate visual sensory sampling (98), suggesting that LGN functional connectivity with other nodes of the allostatic-interoceptive system might reflect polysynaptic connections that are functionally meaningful. In our study, the observation of a broad allostatic-interoceptive system is consistent with the confirmed monosynaptic connections between the *a priori* ROIs and the understanding that functional connectivity may reflect both monosynaptic and polysynaptic connections (99).

The connecting ‘hub’ regions of the allostatic-interoceptive system observed at 7 Tesla covered *all* hypothesized cortical regions of interest, including the full extent of primary interoceptive cortex (dpIns, dmIns; (15)) and the primary visceromotor regions (vAIns, sgACC, pACC and aMCC; (100)). Several other connecting ‘hub’ regions (MCC, PCC, IFG, PHG, STG) were also observed and we confirmed their anatomical connections to documented allostatic regions in non-human animals. The remaining connecting regions (i.e., MFG, SFG, isthmus of the cingulate, cuneus) have not been documented as having monosynaptic anatomical connections to our subcortical and cortical seed regions – their functional connectivity may reflect polysynaptic connections or novel connections in humans. Importantly, most of the additional connecting regions observed at 7 Tesla (i.e., pACC, PCC, isthmus cingulate, SFG, MFG and mIns; except the sgACC) belong to the ‘rich club’ (the most densely interconnected regions in the cortex and thought to serve as the “backbone” that synchronizes neural communication throughout the brain; (105)), consistent with the hypothesized central role of the allostatic-interoceptive system as a high-capacity “backbone” for integrating information across the entire brain (106).

The results of this study have several important functional implications. First, a number of brain regions within the allostatic-interoceptive network most likely play an important role in coordinating and regulating the systems of the body even though they are involved in other psychological phenomena. For example, the SC is typically studied for visuomotor functioning in humans but has been shown to be important for approach and avoidance behavior as well as the accompanying changes in visceromotor activity in non-human mammals (e.g., 62, 63, 107) via anatomical connections to ACC (50) and hypothalamus (51). Similarly, the hippocampus is usually considered central to memory function, but evidence from non-human animals indicates that the hippocampus also plays a role in the regulation of feeding behaviors and in interoception-related reward signals (108–113). There is also circumstantial evidence that interoceptive signals, relayed from the vagus nerve to the hippocampus via the NTS and septal nuclei, may play a role in event segmentation (114, 115). Furthermore, the LGN is usually considered part of the visual pathway that relays visual information from the retina and the cerebral cortex. However, the current functional connectivity findings are consistent with tract-tracing evidence showing LGN’s monosynaptic connections with cortical (e.g., pACC (69)) and subcortical visceromotor structures (e.g., hypothalamus (68), PAG (52), and PBN (116)), suggesting a role for facilitating communication among brain structures implicated in bodily regulation, in addition to its role in integrating interoceptive and visual signals (39). The broad functional connectivity profile of the LGN is also consistent with evidence of tracts between the LGN and other hypothesized regions of the allostatic-interoceptive network such as the hippocampus, amygdala, DR, SC, SN, and LC (**Supplementary Table 1**).

Second, both the default mode and salience networks have been functionally implicated in cardiovascular regulation as well as other aspects of allostasis (9, 117, 118), and they have also been implicated in mental and physical illness and their comorbidities (e.g., (119, 120)). Not surprisingly, psychiatric illnesses (e.g., depression (121, 122), schizophrenia (123, 124)), neurodevelopmental illnesses (e.g., sensory processing disorder/autism spectrum disorder (125, 126)), neurodegenerative illnesses (e.g., dementia/Alzheimer’s disease (127, 128), Parkinson’s disease (129, 130)) and physical illnesses (e.g., heart disease (131), chronic pain (132)) present with symptoms related to altered interoception or visceromotor control, and some of these symptoms are transdiagnostic (133–135). Moreover, interoceptive and visceromotor symptomatology is often accompanied by altered neurobiology (e.g., volume, structural connectivity, functional connectivity, evoked potential, task activation) in the allostatic-interoceptive system (e.g., depression: (136, 137); autism: (138); dementia: (33, 127, 128); chronic pain: (139); transdiagnostic: (140–143)). In addition, there is evidence showing that psychological therapies targeting interoceptive processes (144) and neuromodulations targeting distributed regions within the allostatic-interoceptive system (145, 146) may be effective transdiagnostic interventions. Taken together, these findings suggest that altered function of the allostatic-interoceptive system may be a transdiagnostic feature of mental and physical illness that holds promising clinical utility. More fundamentally, the system identified in this paper provides a scientific tool for integrating studies across psychological and illness domains in a manner that will speed discovery, the accumulation of knowledge and, potentially, strategies for more effective treatments and prevention.

Finally, the findings reported here are consistent with the growing body of evidence that a number of subcortical and cortical brain regions are important during both the regulation of bodily functions and during cognitive phenomena, calling into question their functional segregation (147–149). Our findings suggest that the default mode and salience networks may be concurrently coordinating, regulating and representing organs and tissues of the internal milieu at the same time that they are engaged in a wide range of tasks spanning cognitive, perceptual, emotion and action domains (see Figure 5 in (19)) (38, 150–154). Therefore, our results, when situated in the broader published literature, suggest that the default mode and salience networks create a highly connected functional ensemble for integrating information across the brain, with interoceptive and allostatic signaling at the core. Regulation of the body has been largely ignored in the neuroscientific study of the mind, in part because much of interoceptive modeling occurs outside of human awareness (18, 134).

Several limitations within the current study should be addressed in future studies. First, we did not validate the connectivity strength within the allostatic-interoceptive network against signal-based measures of interoception (e.g., heart-beat evoked potentials), although there is growing evidence that even at rest, limbic regions of the brain continually issue allostatic control signals and there should be synchronous relationships between resting state BOLD signals and electrical signals from visceromotor movements (155). Second, we did not fully monitor participants’ wakefulness (e.g., via video recording) during the three ten-minute resting state scans. The default mode and salience networks are present during sleep (156), although the strength of within-network functional connectivity has been shown to vary (with evidence of both stronger and weaker connectivity) as a function of wakefulness (157–160). Third, we did not map every relevant subcortical area that may be involved in allostasis or interoception. For example, opportunities for further research include septal nuclei (with direct projections to limbic regions such as the hippocampus and implicated in temporal control of neurons that make up the allostatic-interoceptive network; (161, 162)), circumventricular organs (e.g., area postrema with unique access to peripheral signaling molecules via its permeable blood-brain barrier; (163, 164)), and motor brainstem nuclei (e.g., dorsal motor nucleus of the vagus and nucleus ambiguus whose neurons give rise to the efferent vagus nerve; (165, 166)). As several of the known anatomical connections of the hypothalamus were not observed, further investigation of hypothalamic functional connectivity is warranted. Further investigation should use different hypothalamic nuclei or subregions as seeds may given functional heterogeneity in relation to allostatic processes (167, 168). The cerebellum is also likely involved in allostasis and interoception (169, 170).

## Online Methods

### Participants and MRI acquisition

We recruited 140 native English-speaking adults, with normal or corrected-to-normal vision, and no history of neurological or psychiatric conditions. All participants provided written informed consent in accordance with the guidelines set by the institutional review board of Massachusetts General Hospital. Forty-nine participants were excluded from the current analysis (19 withdrew prior to the MRI session, three withdrew during MRI acquisition due to discomfort, six did not complete scans due to online scan reconstruction failure, three did not complete scans due to time constraint, four were excluded due to other technical issues during acquisition, 10 were excluded due to scanner sequence error, four were excluded due to corrupted MRI data that could not be processed and one was excluded due to excessive artifacts in the structural scan). This resulted in a final sample of 90 participants (26.9 ± 6.2 years old; 40 females, 50 males). MRI data were acquired using a 7 Tesla scanner (Magnetom, Siemens Healthineers, Erlangen, Germany) with a 32-channel phased-array head coil and personalized padding to achieve a tight fit. Participants completed a structural scan, three resting state scans of 10 minutes each, three diffusion-weighted scans, as well as other tasks unrelated to the current analysis. At the beginning of each resting state scan, participants were instructed to keep their eyes open and indicated their readiness to start the scan via button press. MRI parameters are detailed in SI.

### Preprocessing of fMRI data

The preprocessing pipeline began with reorientation, slice timing correction, concatenation of all three resting state runs, coregistration to the structural T_1_-weighted image, and motion correction (framewise displacement mean = 0.17, s.d. = 0.14, with 98.7% of frames showing sub-voxel motion; (171)). We then conducted nuisance regression to remove physiological noise due to motion (six parameters measuring rotation and translation), as well as due to non-BOLD effects evaluated in the white matter, ventricular cerebrospinal fluid, and the cerebral aqueduct. We then conducted temporal filtering and normalization. Finally, we performed conversion to Freesurfer orientation/dimensions, detrending, spatial smoothing (1.25mm), and resampling to cortical surfaces. Preprocessing details are provided in SI.

### Functional Connectivity Analysis

Seven cortical seeds (4mm-radius spheres) were defined based on previous fMRI studies of interoception using the procedure outlined in (172). The 14 subcortical seeds were defined based on the Brainstem Navigator toolkit (https://www.nitrc.org/projects/brainstemnavig/) (e.g., (95)), CANLAB Combined Atlas 2018 (github.com/canlab), and Freesurfer segmentation (e.g., (173)). See **Supplementary Methods** for details about seed definition. We randomly resampled 80% of the sample (*N* = 72) 1000 times. In each iteration, for each seed, we estimated cortical connectivity using Freesurfer-based analysis procedure as outlined in (19). This yielded final group maps that showed regions whose fluctuations significantly correlated with the seed’s fMRI time series, which were binarized to retain positive connectivity surviving the threshold of *p* < .05 and summed across 1000 iterations to obtain ‘bootstrapped connectivity’ maps. We also quantified seed-to-seed functional connectivity by computing Pearson’s correlation coefficient between all pairs of ROIs and applying the Fisher’s *r*-to-*z* transform. Significance at the group level was assessed with a two-tailed one-sample t test.

### Connecting Regions and K-Means Cluster Analysis

To visualize the connecting ‘hub’ regions, we conjoined binarized functional connectivity maps (*p* < 0.05) for all seeds. To replicate the previously discovered two-subsystem distinction within the allostatic-interoceptive network (19), we first computed a similarity matrix capturing pairwise *η^2^* (174) between the un-thresholded bootstrapped group maps of cortical seeds and then applied k-means clustering algorithm (*kmeans*, MATLAB) with a range of *k* between 2-10 (for each *k,* we tested 10 initializations with new centroid positions, each with a maximum of 1000 iterations to find the lowest local minimum for sum of distances). We evaluated the optimal *k* using the Calinski-Harabasz Criterion (79). To visualize each subsystem, we binarized the group connectivity maps (*p* < 0.05) and calculated the conjunction between maps within the same cluster.

### Data and Code Accessibility

Raw and preprocessed data can be found at: https://openneuro.org/datasets/ds005747. Analysis outputs and codes can be found at: https://github.com/jiahez/7-Tesla-Allostatic-Interoceptive-System.

## Acknowledgments

This work was supported by grants from the National Institutes of Health (NCI U01 CA193632, R01 AG071173, R01 MH109464, R01 MH113234, NIDCD R21 DC015888, NIBIB K01 EB019474, and NIA R01 AG063982), the National Science Foundation (BCS 1947972), the U.S. Army Research Institute for the Behavioral and Social Sciences (W911NF-16-1-019), and the Unlikely Collaborators Foundation. The views, opinions, and/or findings contained in this review are those of the authors and shall not be construed as an official Department of the Army position, policy, or decision, unless so designated by other documents, nor do they necessarily reflect the views of the Unlikely Collaborators Foundation.

## Author contributions

T.W., L.W., A.B.S., L.F.B. and M.B. designed research. J.Z., D.C., J. T., L.H., K.M.L, K.M., A.B.S., K.S.Q., S.W-G., L.F.B. and M.B. performed research. J.Z., D.C., P.D., T.S., L.F.B. and M.B. analyzed data and wrote the paper. All authors read and approved the paper. Competing interest statement: The authors declare no conflict of interest.

