## Supplement for "Cortical and subcortical mapping of the allostatic-interoceptive system in the human brain using 7 Tesla fMRI"

**Supplementary Table 1. Correspondence between functional connectivity observed in the current study and tract-tracing results in non-human animals, demonstrating anatomical connections between cortical and subcortical ROIs.**

| | | **Cortical regions** | | | | |
|---|---|---|---|---|---|---|
| | | **sgACC (BA25)** | **pACC (BA24, 32)** | **aMCC (BA24)** | **mvaIns/lvaIns** | **dmIns/dpIns** |
| **Cortical regions** | **sgACC (BA25)** | | | | | |
| | **pACC (BA24, 32)** | *†Fig. 1* (1)<br>†Fig. 5 (2)<br>†Fig. 2A (1) | | | | |
| | **aMCC (BA24)** | *†Fig. 4* (3)<br>†Fig. 7 (2)<br>†Fig. 3A (1) | *†Fig. 5* (2)<br>*†Fig. 2A* (1)<br>†Fig. 7 (2)<br>†Fig. 3A (1) | | | |
| | **mvaIns/lvaIns** | *†Fig. 8* (4)<br>†Case M707 (5) | *†Fig. 5* (2)<br>†Case M776 (5) | *†Fig. 6* (2)<br>*†Fig. 1* (6)<br>†Fig. 1 (7) | | |
| | **dmIns/dpIns** | *†Not observed* (1, 6, 8–10)<br>†Not observed (5, 6, 8, 9) | *†Fig. 5* (2)<br>†Not observed (5, 6, 8, 9) | *†Fig. 3* (6)<br>†Fig. 4 (7) | *†Fig. 1* (7)<br>†Fig. 4 (7)<br>†Fig. 1 (6) | |

Note: Gray shading indicates connections that showed significant positive functional connectivity in the current study. Text describes the results of tract-tracing studies of non-human animals. Efferent connections (i.e., from the ROI in the column to the ROI in the row) are indicated in regular font, afferent connections (i.e., from the ROI in the row to the ROI in the column) are indicated in italic font. We considered two regions to share anatomical connections if the anatomical tracer injected in one region showed any terminations in the second region. * rat study, † monkey study, ʃ mouse study, ˆ cat study, " hamster study, ** rabbit study

**Supplementary Table 1 (continued). Correspondence between functional connectivity observed in the current study and tract-tracing results in non-human animals, demonstrating anatomical connections between cortical and subcortical ROIs.**

| | | **Cortical regions** | | | | |
|---|---|---|---|---|---|---|
| | | **sgACC (BA25)** | **pACC (BA24, 32)** | **aMCC (BA24)** | **mvaIns/lvaIns** | **dmIns/dpIns** |
| **Subcortical regions** | **mdThal** | ^Fig 3 (11)<br>†Fig 4 (12) | ^Fig 3 (11)<br>*†Fig 19(13)* | Not evident | **Fig 6 (14)*<br>*Case R579 (15)<br>*Fig 3, 4, Table 1 (16) (bidirectional) | **Fig 6A(17)* |
| | **LGN** | Not evident | *"Fig 7 (18)* | Not evident | *Not observed (6) | *Not observed (6) |
| | **Hypothal** | †Fig 4,10 (19)<br>†Fig 8 (20)<br>†Fig 2 (12) | **Fig 3 (21)*<br>†Fig 4,9,12 (19)<br>†Fig 9 (20)<br>^Fig 3 (11) | †Fig 4 (19) | *Fig 3,5 (16)<br>*Fig 2 (21)<br>†Fig 3,6 (19)<br>(bidirectional) | *Fig 2 (21)<br>†Fig 6 (19) |
| | **Hippo** | *†Fig 6 (22)*<br>*†Fig 3 (23)* | *†Fig 4 (22)* | *†Fig 5,7 (22)*<br>*†pg 440 (23)* | **Fig 12 (24)* | **Not observed (24)* |
| | **Amygdala** | †Fig 2,5,6 (25) (bidirectional)<br>†Fig 3,4 (26)<br>†Fig 2,5,6 (27) | ^Fig 3 (11)<br>†Fig 2,6 (25) (bidirectional)<br>†Fig 3,4 (26)<br>†Fig 2 (27) | †Fig 2,5,6 (25) (bidirectional)<br>†Fig 3,4,5 (26)<br>†Fig 2,7 (27) | †Fig 2 (28)<br>†Fig 2 (29)<br>†Fig 3,4 (26)<br>†Fig 2 (27) | †Fig 3 (28)<br>†Fig 2,3,6 (29)<br>†Fig 3,4,5 (26)<br>†Fig 2,8,9 (27) |
| | **Striatum** | ^Fig 3 (11)<br>†Fig 3-5 (30) (undirectional) | ^Fig 3 (11)<br>^Fig 3 (11)<br>†Fig 3-5 (30) (undirectional) | †Fig 3-5 (30) (undirectional) | †Fig 3-15 (31)<br>†Fig 4,6,7,9 (32) | †Fig 3-15 (31)<br>†Fig 4,6-9,11 (32) |
| | **PAG** | †Fig 2-4 (12)<br>†Fig 1-6 (33) | ^Fig 3 (11)<br>†Fig 1-6 (33) | †Fig 1-7 (33) | †Fig 1,7 (33)<br>*Fig 3,5 (16) (bidirectional) | †Fig 7 (33)<br>*Fig 7 (16) |
| | **DR** | †Fig 8 (12) | †Fig 12 (34)<br>*Fig 8 (35) | †Fig 12 (34) | *Fig 2 (35)<br>*Fig3,5 (16) (bidirectional) | *Not observed (35) |
| | **SC** | *Fig 15 (36) | *Fig 15 (36) | *Fig 15 (36) | *Fig 5 (37) | Not evident |
| | **SN** | *pg 1757 (36) | *pg 1757 (36) | *Not observed (36) | *Fig 3 (16) | Not evident |
| | **VTA** | *pg 1757 (36) | *pg 1757 (36) | *Not observed (36) | *Fig 3 (16) | Not evident |
| | **PBN** | †Fig 8 (12) | †Fig 12 (34)<br>^Fig 3 (11) | †Fig 12 (34) | *Fig 1,3 (38) (bidirectional)<br>*Fig 3,5[10] (bidirectional) | **Fig 4 (14)*<br>*Fig 1,3 (38) (bidirectional) |
| | **LC** | Not evident | †Fig 12 (34)<br>**Table 1 (39)* | †Fig 12 (34) | *Fig 5 (16) | Not evident |
| | **NTS** | †Not observed (12)<br>*Not observed (40) | *Not observed (40)<br>*Fig 1 (41) | *Not observed (40) | *Fig 4 (40) | *Fig 5 (42) |

Note: Gray shading indicates connections that showed significant positive functional connectivity in the current study. Text describes the results of tract-tracing studies of non-human animals. Efferent connections (i.e., from cortical to subcortical) are indicated in regular font, afferent connections (i.e., from subcortical to cortical) are indicated in italic font. We considered two regions to share anatomical connections if the anatomical tracer injected in one region showed any terminations in the second region. We reviewed the entire striatum because many source studies did not differentiate dorsal vs. ventral portions of the striatum. * rat study, † monkey study, ʃ mouse study, ^ cat study, " hamster study, ** rabbit study

**Supplementary Table 1 (continued). Correspondence between functional connectivity observed in the current study and tract-tracing results in non-human animals, demonstrating anatomical connections between cortical and subcortical ROIs.**

| | | **Subcortical regions** | | | | | | | | | | | | | |
|---|---|---|---|---|---|---|---|---|---|---|---|---|---|---|---|
| | | **mdThal** | **LGN** | **Hypothal** | **Hippo** | **Amygdala** | **Striatum** | **PAG** | **DR** | **SC** | **SN** | **VTA** | **PBN** | **LC** | **NTS** |
| **Subcortical regions** | **mdThal** | | | | | | | | | | | | | | |
| | **LGN** | Not evident | | | | | | | | | | | | | |
| | **Hypothal** | _*Fig 24 (43)_ | *Fig 2 (44)<br>*Fig 9 (45)<br>*Fig 3 (46) | | | | | | | | | | | | |
| | **Hippo** | Not evident | *Table 6 (47) | _*Fig 1 (48)_<br>_*Fig 3 (47)_ | | | | | | | | | | | |
| | **Amygdala** | _*Fig 24 (43)_<br>_*Fig 5 (49)_ | *Fig 3 (48) | _*Fig 1(50)_<br>_*Fig 1 (51)_ | *Fig 5 (52) | | | | | | | | | | |
| | **Striatum** | _*Fig 22 (43)_ | Not evident | _*Fig 2,3_ (53)<br>_*Fig 1_ (15) | †Fig 5 (54) | †Fig 4 (54)<br>*Fig 2,3 (53) | | | | | | | | | |
| | **PAG** | _†Case SM-7,8_ (55)<br>*Fig 2 (56)<br>_*Fig 6_ (57) | _*Case R-43, 45, 59_ (58) | †Case SM-8 (55)<br>*Fig 9, 10 (57)<br>_*Case R-43, 45, 50, 59_ (58) | *Case 33 (59) | _*Case R-50,R-59_ (58)<br>^Case H3, 7, 19 (60) | Not evident | | | | | | | | |
| | **DR** | _*Fig 2_ (61) | _*Fig 3_ (62)<br>_^Fig 1_(63)<br>_*Fig 3_ (64) | _*Fig 3_ (65)<br>_*Fig 9_ (61) | _*Fig 2_ (66) | _*Fig 2_ (61) | _*Fig 9, 13_ (61) | _*Case R-49, 50, 59_ (58) | | | | | | | |
| | **SC** | *Fig 10 _(43)_<br>*Not observed (67) | *Fig 3 (64)<br>_† Fig 1_(68)<br>*Fig 3 (69)<br>_*R27_ (58) | _^Fig 3_ (70)<br>_**Fig 1_ (71) | *Not observed (72) | _*Fig 4,6_ (73)<br>_*pg 403_ (74) | Not evident | ^Fig 1(75) (bidirectional)<br>*Fig 2 (76) | *Table 2 (77)<br>*Fig 1 (62) | | | | | | |
| | **SN** | _*Fig 26 (43)_<br>^Fig 4 (78) | _^Fig 1_ (63) | _*Fig 6_ (79) | _*pg 9_ (80) | _*Fig 1_(81)<br>_*Fig 1_ (82)<br>_*Fig 6_ (79) | _*Fig 2_ (83)<br>_^Fig 4_ (84)<br>_*Fig 3_ (85)<br>_*Fig 6_ (79) | _*Fig 2_ (86)<br>**Fig 4 (87) | *Fig 3 (88)<br>*Fig 2 (61) | ^Fig 2 (89)<br>^Fig 2 (60)<br>*Fig 3 (90) | | | | | |
| | **VTA** | _*Fig 31(43)_<br>^Fig 3 (78) | Not evident | _*Fig 4_ (72) | *Not observed (72) | _*Fig 12_ (72) | _*Fig 12_ (72) | _*Fig 3_ (86)<br>**Fig 4 (87) | *Fig 2 (61)<br>*Fig 6 (65) | _^Fig 6_ (91)<br>*Fig 1 (90)<br>^Fig 1 (89) | _*Fig 12_ (72) | | | | |
| | **PBN** | _*Fig 9R (43)_ | _^Fig 1_ (92)<br>^Fig 1 (63) | _"Fig 3_ (93)<br>_*Fig 2_ (94) | _*Fig 2_ (95) | _"Fig 2_ (93)<br>_*Fig 1_ (96) | _*Fig 2_ (97) | _*Case R-43, 45, 50, 59_ (58) | _*Fig 6_ (65) | _*Fig 3_ (69)<br>_*Fig 3_ (98) | _*Fig 8_ (83)<br>*Fig 4 (79) | *Fig 2 (99)<br>*Fig 18 (72) | | | |
| | **LC** | _*Fig 9R (43)_<br>^Fig 5 (78) | _*Fig 8_ (100)<br>_*Fig 10_ (64) | *Table 1 (101)<br>*Fig 23 (102)<br>_*Fig 3_ (103) | _*Fig 1C_ (104)<br>_*Fig 2_ (95)<br>_*Fig 2_ (105)<br>_*Fig 3_ (103) | *Table 1C (101) | Not evident | _*Case R-43, 45, 50_ (58) | *Fig 15 (102)<br>_*Fig 3_ (106)<br>_(bidirectional)_ | _*Fig 4,5_ (77) | *Not observed (79) | *Fig 1 (107) | *Fig 17 (102) | | |
| | **NTS** | _*Fig 31 (43)_ | Not evident | _*Fig 2_ (108)<br>ʃFig 1 (109) | *Not observed (95) | _*Fig 4_ (108)<br>_*Fig 5_ (110) | *Fig 1 (111)<br>*Fig 1 (112)<br>*Fig 4 (113) | _*Fig 3_ (86)<br>**Fig 4 (87) | _*Fig 2_ (114) | _*Fig 4_ (40)<br>*Not observed (113) | Not evident | _*Fig 2_ (115)<br>_*Table 2_ (116)<br>*Fig 1 (111) | "Fig 3 (93)<br>*Fig 1(113)<br>ʃFig 1 (109) | _*Fig 1_ (117)<br>_*Not observed_ (102) | |

Note: Gray shading indicates connections that showed significant positive functional connectivity in the current study. Text describes the results of tract-tracing studies of non-human animals. Efferent connections (i.e., from the ROI in the column to the ROI in the row) are indicated in regular font, afferent connections (i.e., from the ROI in the row to the ROI in the column) are indicated in italic font. We considered two regions to share anatomical connections if the anatomical tracer injected in one region showed any terminations in the second region. We reviewed the entire striatum because many source studies did not differentiate dorsal vs. ventral portions of the striatum. * rat study, † monkey study, ʃ mouse study, ^ cat study, "hamster study, ** rabbit study. The upper triangle has been grayed out.

**Supplementary Table 2.** Functional connectivity of the superior parietal lobule (averaged Fisher's transformed *z*-scores). The seed coordinates of the SPL are MNI X, Y, Z = 46, 12, 29 (118).

| ROI | Functional connectivity |
|---|---|
| mdThal | |
| LGN | 0.02 |
| Hypothal | |
| Hippo | 0.12* |
| dAmy | 2.50* |
| NAcc | |
| PAG | |
| DR | |
| SC | 0.09* |
| SN | 0.02 |
| VTA | |
| PBN | |
| LC | |
| VSM | 0.03* |

Note. Empty cells indicate negative connectivity. $*p < 0.05$

**Supplementary Table 3.** Percentage of subcortical ROI that overlapped with the subcortical connecting regions. Subcortical connecting regions were computed by taking the conjunction between the binarized maps ($p < 0.05$) of all subcortically seeded functional connectivity maps ($N = 14$).

| ROI | Percentage |
|---|---|
| mdThal | 1.1 |
| LGN | 26.8 |
| Hypothal | 0 |
| Hippo | 1.8 |
| dAmy | 4.2 |
| NAcc | 0.2 |
| PAG | 12.0 |
| DR | 14.4 |
| SC | 11.4 |
| SN | 12.5 |
| VTA | 9.1 |
| PBN | 0 |
| LC | 0 |
| VSM | 0 |

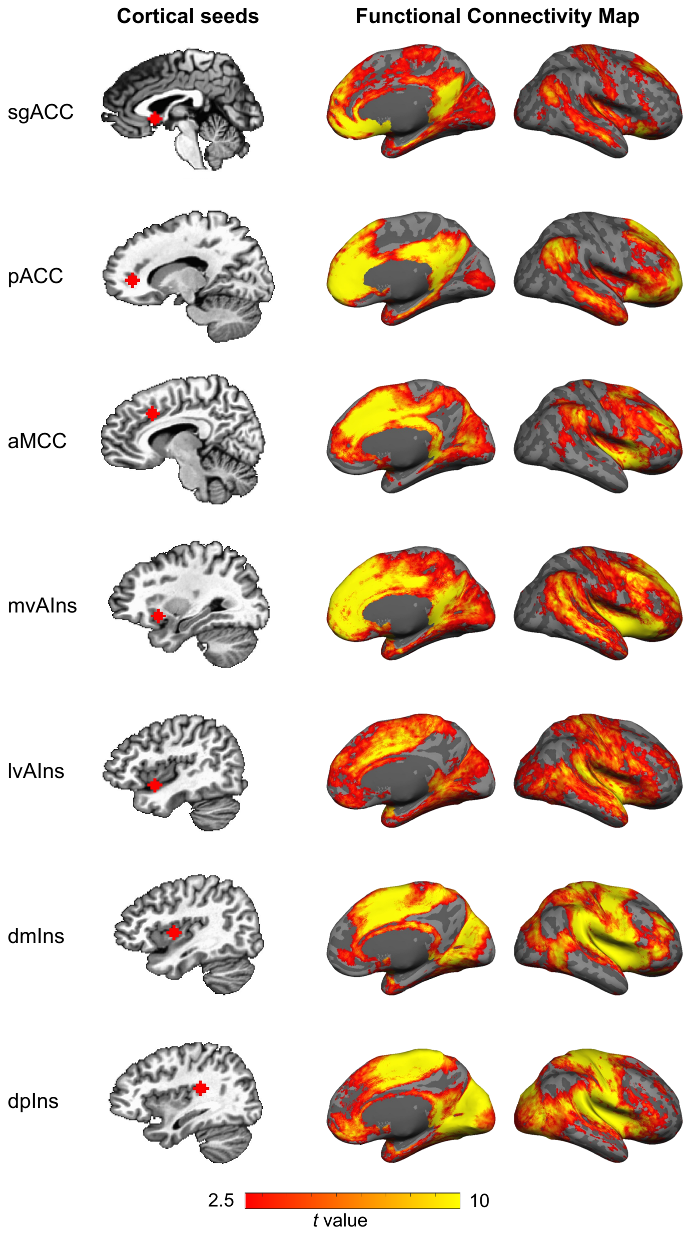


**Supplementary Figure 1.** Cortico-cortical functional connectivity displayed as *t* values ($N = 90$). The maps were masked by voxels that showed positive connectivity at a threshold of $p < .05$ in more than 950 iterations of the 1000 subsampled analyses (i.e., bootstrapped connectivity ≥ 950/1000).

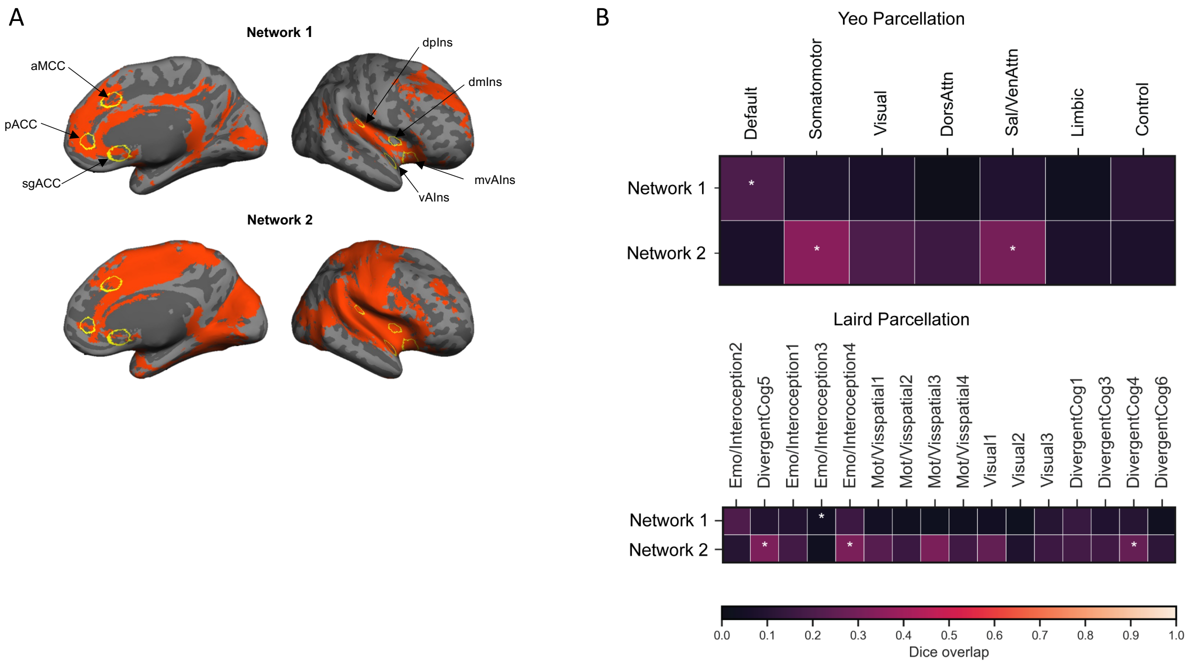


**Supplementary Figure 2.** The two large-scale intrinsic networks composing the cortical allostatic-interoceptive system correspond to the default mode network and salience/somatomotor networks. **(A)** The cortical allostatic-interoceptive system is composed of two large-scale intrinsic networks. K-means clustering (*k* = 2, 1000 iterations) yielded the most optimal solution where Network 1 (resembling the default mode network) included a cluster of maps seeded in the sgACC, aMCC, pACC and mvAIns, and Network 2 (resembling the salience network) included a cluster of maps seeded in the lvAIns, dmIns and dpIns. All displayed maps result from the conjunction of binarized maps ($p < 0.05$) in the same cluster. Cortical ROIs are outlined in yellow (ROI names are labeled in the top panel). **(B)** We computed Dice overlap between network maps and the Yeo 7-network cortical parcellation (119) using the Network Correspondence Toolbox (https://github.com/rubykong/cbig_network_correspondence) (120). In the grids, cells with significant Dice overlap at $p < .05$ (i.e., showing substantial correspondence) are denoted with an asterisk. Network 1 showed significant Dice overlap solely with the default mode network, while Network 2 showed significant Dice overlap with the salience/ventral attention network and somatomotor network.

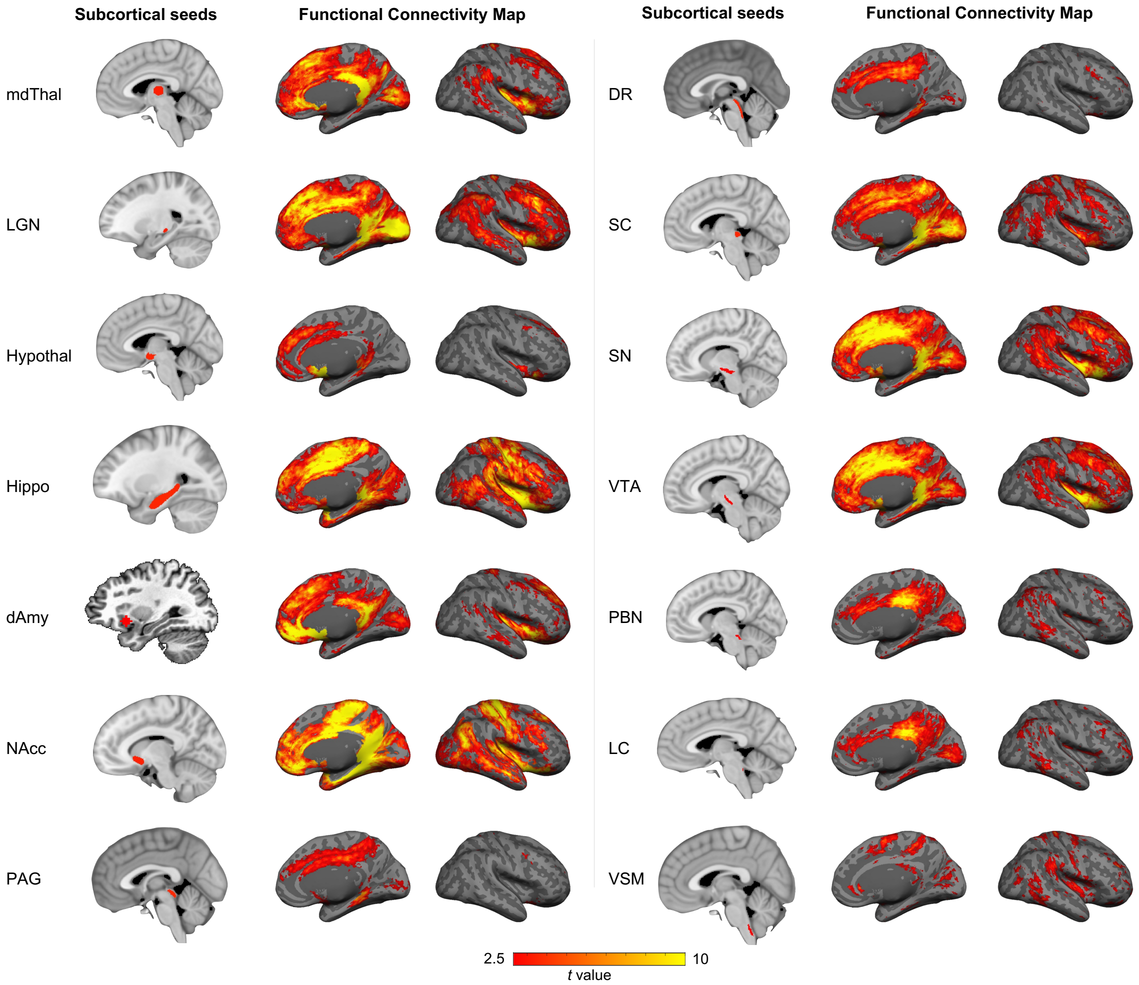


**Supplementary Figure 3.** Subcortico-cortical functional connectivity displayed as *t* values ($N = 90$). The maps were masked by voxels that showed positive connectivity at a threshold of $p < .05$ in more than 950 iterations of the 1000 subsampled analyses (i.e., bootstrapped connectivity ≥ 950/1000).

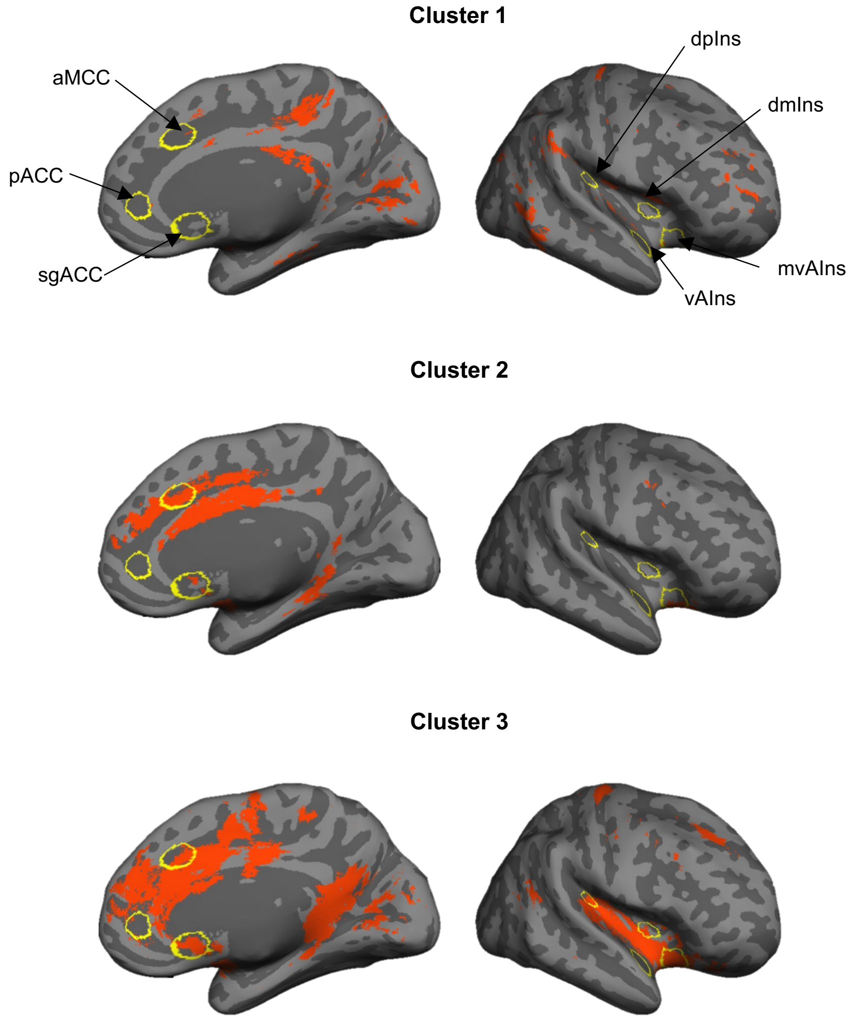


**Supplementary Figure 4.** Clustering solution (*k* = 3) for cortical maps of subcortical allostatic-interoceptive seeds. Cluster 1 included maps that were seeded in small lower brainstem ROIs (LC, PBN, VSM). Cluser 2 included maps that were seeded in small upper brain stem ROIs (PAG and DR) and the hypothalamus. Cluster 3 included maps that were seeded in larger subcortical seeds (mdThal, LGN, hippocampus, dAmy, NAcc, SC, SN and VTA). All displayed maps result from the conjunction of binarized maps ($p < 0.05$) in the same cluster. Cortical ROIs are outlined in yellow (ROI names are labeled in the top panel).

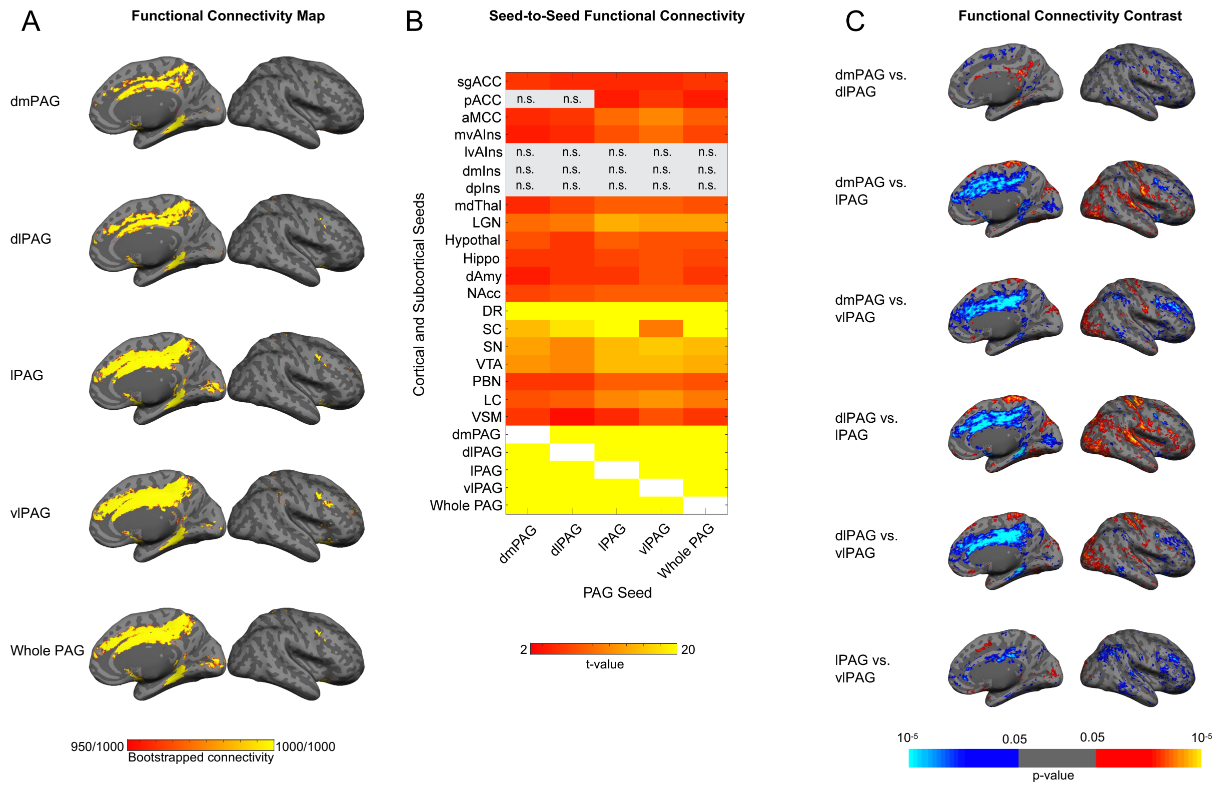


**Supplementary Figure 5.** Intrinsic connectivity of PAG and its subregions within the allostatic-interoceptive system. Panel **(A)** shows bootstrapped connectivity maps obtained from resampling 80% of the sample ($N = 72$) 1000 times. Panel **(B)** shows connectivity strength between PAG seeds and all other seeds ($p < .05$, uncorrected; white color indicates correlation =1 and gray color indicates subthreshold correlations; $N = 90$). Panel **(C)** shows contrasts obtained by paired-sample t-tests between subregional maps ($p < .05$, uncorrected). lPAG and vlPAG showed more robust and more extensive connectivity than dmPAG and dlPAG, with stronger connectivity especially with aMCC, mvAIns, and dAmy. Abbreviations: dmPAG: dorsomedial PAG; dlPAG: dorsolateral PAG; lPAG: lateral PAG; vlPAG: ventrolateral PAG.

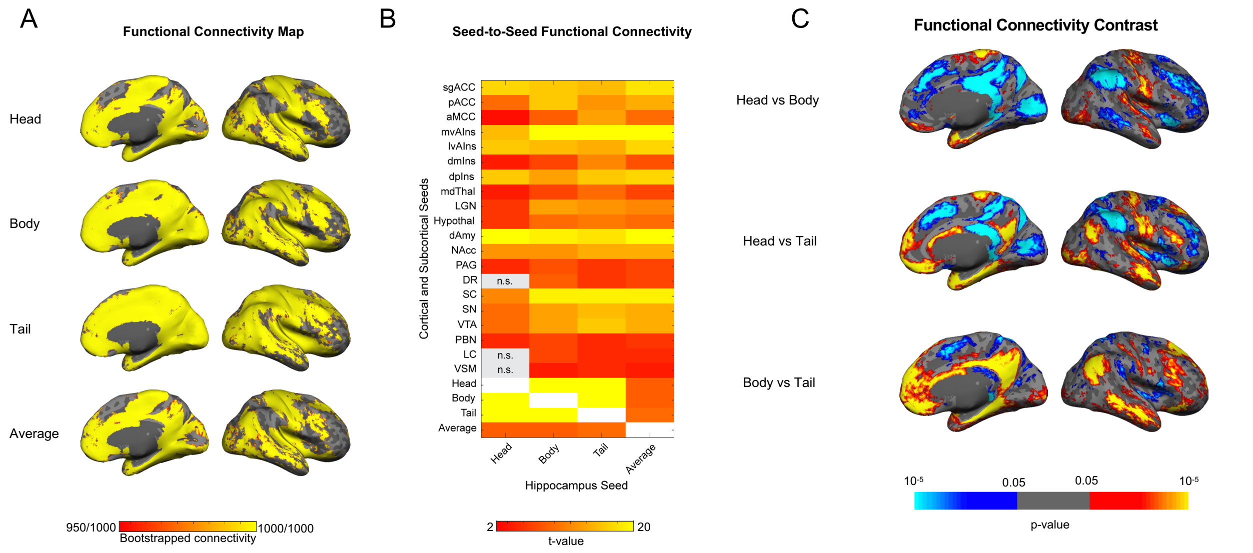


**Supplementary Figure 6.** Intrinsic connectivity of the hippocampus and its subregions within the allostatic-interoceptive system. Panel **(A)** shows bootstrapped connectivity maps obtained from resampling 80% of the sample ($N = 72$) 1000 times. Panel **(B)** shows seed-to-seed connectivity strength between hippocampal seeds and all other seeds ($p < .05$, uncorrected; white color indicates correlation =1 and gray color indicates subthreshold correlations; $N = 90$). Panel **(C)** shows contrasts obtained by paired-sample t-tests between subregional maps ($p < .05$, uncorrected). Hippocampal head and body showed stronger connectivity to default mode nodes such as the MPFC, PCC, AG and lateral temporal cortex. Hippocampal body and tail showed stronger connectivity to salience nodes such as ACC, PCC, SMA, MFG and SMG.

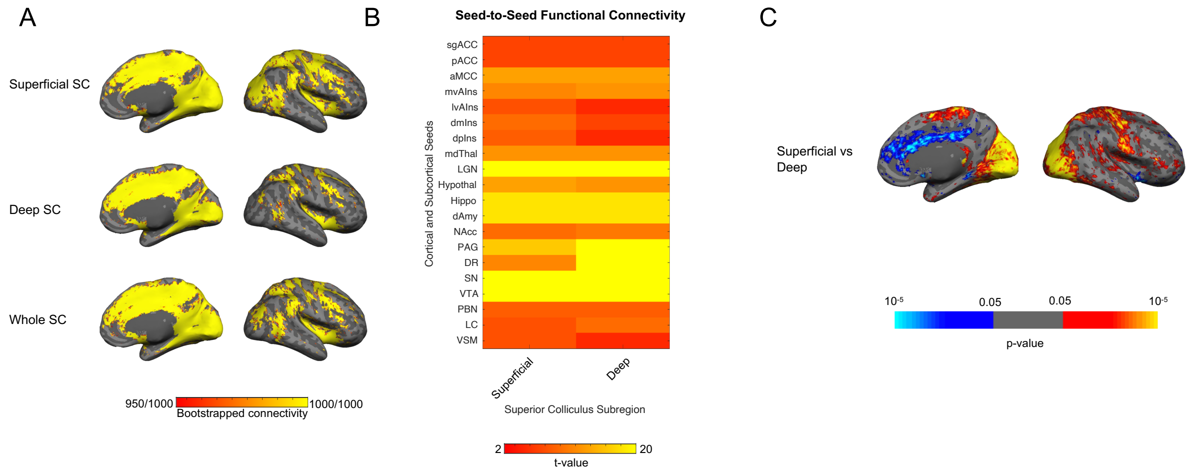


**Supplementary Figure 7.** Intrinsic connectivity of the superficial and deep layers of the SC within the allostatic-interoceptive system. Panel **(A)** shows bootstrapped connectivity maps obtained from resampling 80% of the sample ($N = 72$) 1000 times. Panel **(B)** shows seed-to-seed connectivity strength between SC subregions and all other seeds ($p < .05$, uncorrected; $N = 90$). Panel **(C)** shows contrasts obtained by paired-sample t-tests between subregional maps ($p < .05$, uncorrected). Superficial SC showed stronger connectivity to primary sensory cortices in the posterior insular, occipital, and pericentral regions. Deep SC showed stronger connectivity to allostatic-interoceptive hubs in the vaIns and the entire cingulate cortex.

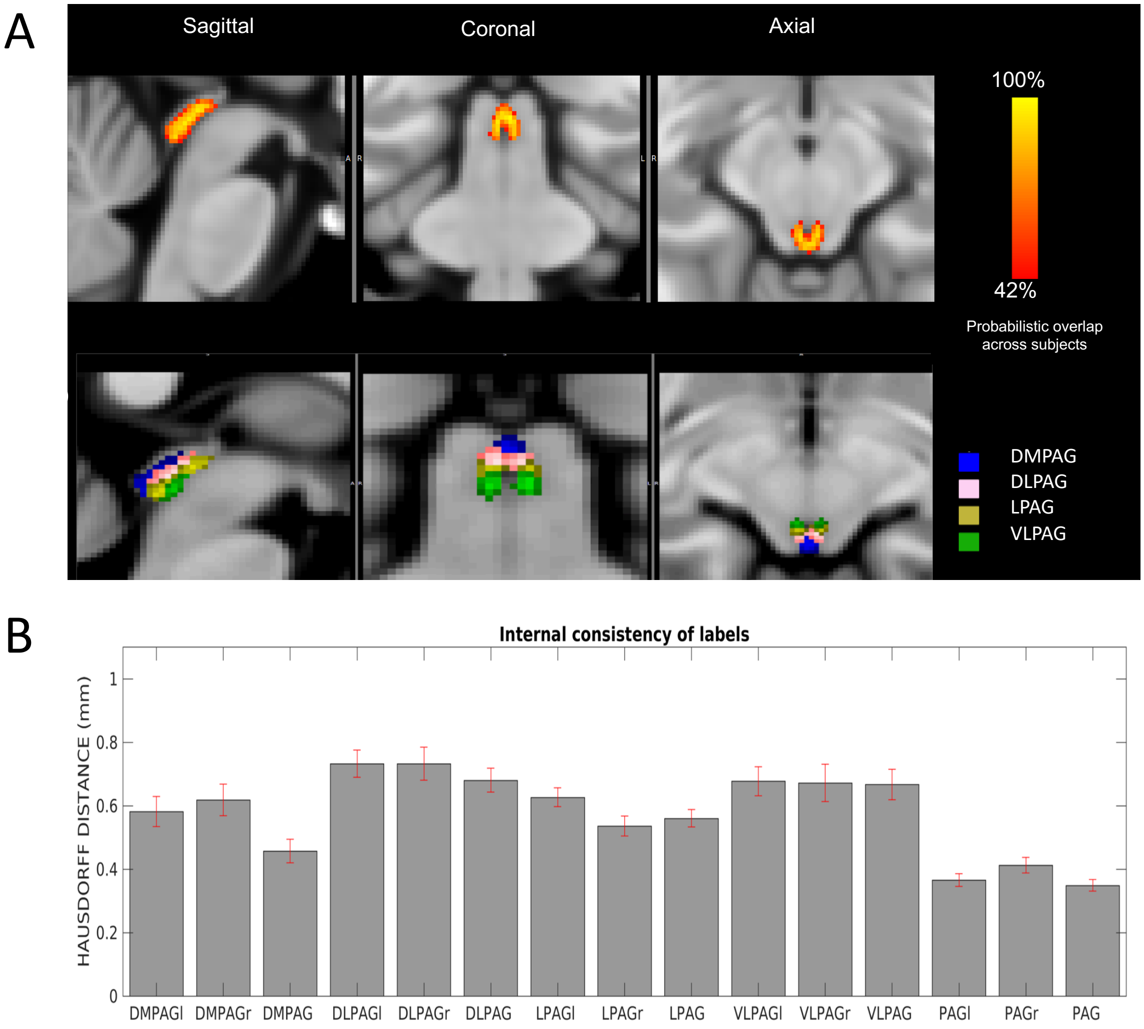


**Supplementary Figure 8. (A)** Probabilistic atlas label of PAG and dorsomedial, dorsolateral, lateral and ventrolateral PAG subregions (DMPAG, DLPAG, LPAG, VLPAG). **(B)** Internal consistency of atlas label, which, for each label, was below the 1.1 mm imaging resolution ($p < 0.05$).

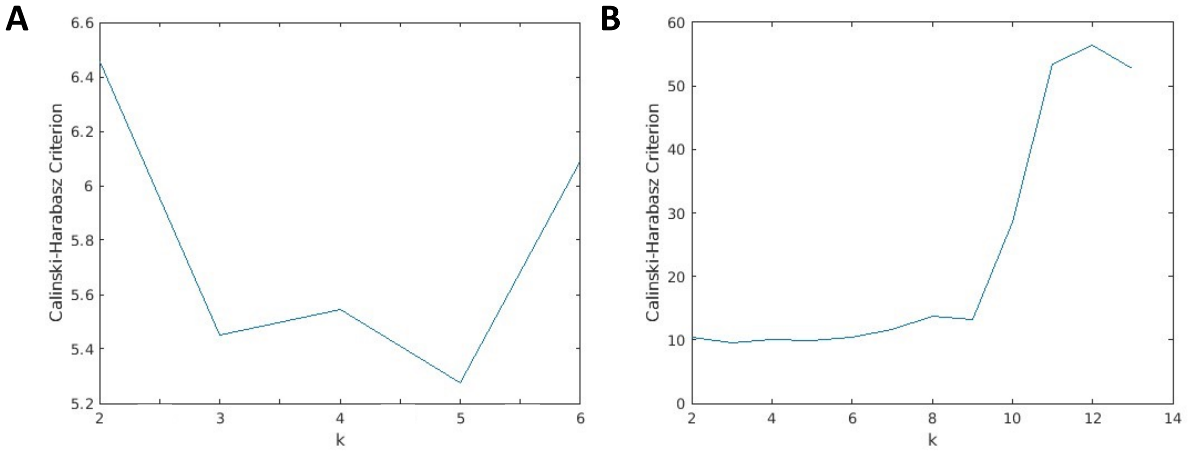


**Supplementary Figure 9.** Evaluation of optimal number of clusters for k-means. We calculated the Calinski-Harabasz Criterion (121) for a range of values of *k* using similarity matrices $\eta^2$ (122) based on cortical maps of **(A)** cortical seeds (for $k$ = 2 to 6) and **(B)** subcortical seeds (for $k$ = 2 to 10). Higher Calinski-Harabasz Criterion indicates larger between-cluster variance and smaller within-cluster variance, i.e., better solution.

**Supplementary Methods**

*MRI acquisition.* MRI data were acquired using a 7 Tesla scanner (Magnetom, Siemens Healthineers, Erlangen, Germany) with a 32-channel phased-array head coil. Participants completed a structural scan, three resting state scans, three diffusion-weighted scans, as well as other tasks unrelated to the current analysis. The structural scan was acquired using a high-resolution multi-echo $T_1$-weighted magnetization-prepared gradient-echo echo-planar image ($T_1$wEPI, (123)) with the following parameters: repetition time (TR) = 8520 ms, echo time (TE) = 22 ms, flip angle (FA) = 90°, voxel size = 1.1 mm isotropic, in plane field of view (FOV) = 205 mm x 205 mm, bandwidth = 1414 Hz/pixel, echo spacing = 0.82 ms, 18 inversion times, GRAPPA-factor = 3. The resting state scans were acquired using a fast low-angle excitation echo-planar technique (1) using the following parameters: TR = 2340 ms, TE = 28 ms, FA = 75°, voxel size = 1.1 mm isotropic, FOV = 205 mm x 205 mm x 135.3 mm, nominal echo-spacing = 0.82 ms, readout bandwidth = 1414 Hz/pixel, N. slices =123, slice acquisition order = interleaved, N. repetitions = 256, phase encoding direction = anterior to posterior, acquisition time = 10'37" per scan, three scans. The diffusion-weighted scans were acquired using a spin-echo echo-planar sequence with parameters: TR = 5800 ms, TE = 63.2 ms, voxel size = 1.1 mm isotropic, N. slices = 61, slice orientation = transversal, readout bandwidth = 1414 Hz/pixel, nominal echo-spacing = 0.82 ms, FOV = 205 mm x 205 mm x 67.1 mm, phase encoding direction = anterior to posterior, GRAPPA-factor = 3, partial Fourier: 6/8, unipolar diffusion-weighting gradients, number of diffusion directions = 60 (b-value ~ 1000 s/mm$^2$), 7 interspersed "b0" images (non-diffusion weighted, b-value ~ 0 s/mm$^2$, also used as $T_2$-weighted MRI), 3 repetitions, acquisition time per repetition 6'58". Seven b0 images were also acquired with the phase encoding direction = posterior to anterior (acquisition time = 1'10").

*Preprocessing of $T_1$-weighted data.* We used FSL (https://fsl.fmrib.ox.ac.uk/fsl/fslwiki), AFNI (https://afni.nimh.nih.gov/), Freesurfer (http://surfer.nmr.mgh.harvard.edu) and SPM (https://www.fil.ion.ucl.ac.uk/spm/) to preprocess the data. For each subject, the $T_1$-weighted EPI ($T_1$wEPI) was first converted to an MPRAGE-like contrast (123), then reoriented to a standard orientation (FSL, reorient2std) and bias field corrected (SPM8, London, UK). The use of $T_1$wEPI has been shown to improve coregistration between structural and functional images, particularly in regions vulnerable to distortion due to magnetic susceptibility differences (124). We then ran standard surface reconstruction and parcellation, as well as subcortical segmentation using Freesurfer.

*Preprocessing of fMRI data.* The preprocessing pipeline for the resting state fMRI scans began with reorientation (FSL, fslreorient2std), slice timing correction (FSL, slicetimer), concatenation of all three resting state runs, coregistration to the structural T1wEPI (FSL, epi_reg using boundary-based registration), and motion correction (Freesurfer, preproc-sess -per-session). We then conducted nuisance regression (custom Matlab script performing general linear model fitting and regression out of covariates of no interest) to remove physiological noise due to motion, as well as due to non-BOLD and pulsatility effects evaluated in the white matter, ventricular cerebrospinal fluid, and the cerebral aqueduct. For the former, we used as nuisance regressors 6 motion parameters (rotations and translations) computed during motion correction. For the latter, we computed the mean time courses (FSL, fslmeants) across voxels of a white matter mask and six cerebrospinal fluid (CSF) masks, which contained voxels in both CSF-spaces distal to the brainstem (such as the lateral ventricle, inferior lateral ventricle, choroid plexus, third ventricle) and neighboring the brainstem (such as the fourth ventricle, and cerebral aqueduct). The cerebral aqueduct mask was manually defined for each subject by computing the signal standard deviation of the resting state time series and selecting the five voxels with the highest standard deviation. Previous work (126, 127) has shown that removal of the signal of the cerebral aqueduct and fourth ventricle is crucial for the functional connectivity analysis of adjacent brainstem nuclei, such as the periaqueductal gray and dorsal raphe. We then conducted temporal filtering (0.01-Hz high-pass filter and 0.08-Hz low-pass filter, AFNI, 3dFourier), and normalization of the fMRI from T1wEPI space to MNI152 space. The latter was performed by computing the affine and non-linear transformation that aligned the $T_1$wEPI image to the 1mm-isotropic resolution $T_1$-weighted MNI152 template (Advanced Normalization Tool, ANTs, Philadelphia, USA), using parameters as in (126). The generic affine transformation was computed by concatenating center-of mass alignment, rigid, similarity and fully affine transformations. The high-dimensional non-linear transformation was a symmetric diffeomorphic normalization transformation with neighborhood cross correlation, regular sampling, gradient step size: 0.15, four multi-resolution levels, smoothing sigmas: 3, 2, 1, 0 voxels – fixed image space –, shrink factors: 6, 4, 2, 1 voxels – fixed image space –, histogram matching of images before registration, data winsorization – quantiles: 0.001, 0.999 –, convergence criterion: slope of the normalized energy profile over the last 10 iterations $< 10^{-8}$. The affine and non-linear transformations were then combined into a single warp field and were applied to the fMRI in T1wEPI space. For each dataset the quality of the coregistration was verified. Finally, we performed conversion to Freesurfer orientation/dimensions (FSL, fslswapdim, fslroi), detrending (FSL, fslmaths), minimal spatial smoothing (1.25mm full-width at half-maximum) (FSL, fslmaths), and resampling to cortical surfaces (Freesurfer, mri_vol2surf).

*Seed definitions.* Seven cortical (sgACC, pACC, aMCC, mvAIns, lvAIns, dmIns and dpIns) and dAmy seeds were defined using the procedure outlined in (118). Cortical seeds were first created using 4mm-radius spheres centered on the MNI coordinates that showed increased activity in previous task-dependent fMRI studies of interoception: dorsal mid insula (dmIns) – 41, 2, 3 (128); dorsal posterior insula (dpIns) – 36, -32, 16 (129); medial ventral anterior insula (mvAIns) – 30, 16, -14 (130); lateral ventral anterior insula (lvAIns) – 44, 6, -15 (131); pregenual anterior cingulate cortex (pACC) – 13, 44, 0 (129); anterior mid cingulate cortex (aMCC) – 9, 22, 33 (131); subgenual anterior cingulate cortex (sgACC) – 2, 14, -6 (132). We found the vertex on the MNI152 pial surface that is closest to each individual subject's cortex and smoothed it by 4 mm. The individual cortical label was projected back into the subject's native volumetric space to calculate the averaged time series within the seed. We directly projected a spherical dorsal amygdala (dAmy) seed – centered on MNI 27, 3, -12 (133) – into each subject's native volumetric space and calculated the averaged time series within the seed. PAG and its four subregions (dorsomedial, dorsolateral, lateral, ventrolateral), were first manually defined in 20 individual subjects based on their diffusion-weighted scans and then the group probabilistic map was thresholded at 35% to generate a group label (**Supplementary Figure 8**; also see section below on *Delineation of the periaqueductal gray and its subregions*). Brainstem nuclei were delineated semi-automatically via 7 Tesla multi-contrast (diffusion and T2-weighted) MRI using neighboring landmarks. Specifically, seven brainstem nuclei seeds were based on the binary labels (35%) from the Brainstem Navigator toolkit (https://www.nitrc.org/projects/brainstemnavig/). Specific Brainstem Navigator labels used for each seed are listed in parentheses following the seed name): 1) locus coeruleus (LC_l, LC_r) (134), 2) lateral geniculate nucleus (LG_r, LG_l) (135), 3) parabrachial nucleus (LPB_l, LPB_r, MPB_l, MPB_r) (136), 4) medullary viscero-sensory-motor nuclei complex (which includes the nucleus tractus solitarius; VSM_l, VSM_r ) (136), 5) dorsal raphe (updated label, see SI; DR) (126), 6) substantia nigra (SN_l, SN_r) (126), and 7) ventral tegmental area (VTA_PBP_l, VTA_PBP_r) (137). The 8$^{th}$ brainstem nucleus, the superior colliculus, and its subregions of superficial and deep layers were defined using a hand-drawn mask template, which was semi-automatically refined based on neighboring landmarks for each subject (138, 139). The medial dorsal thalamus seed was from the medial dorsal thalamus nucleus, defined based on (140), compiled in CANLAB Combined Atlas 2018 (https://github.com/canlab/Neuroimaging_Pattern_Masks/tree/master/Atlases_and_parcellations/2018_Wager_combined_atlas). Hippocampal subregions of head, body, and tail were derived using the longitudinal segmentation method as implemented in Freesurfer (141). The hypothalamic (142) and nucleus accumbens seeds were directly generated from Freesurfer subcortical segmentation (143).

*Preprocessing of diffusion weighted images.* For each subject, diffusion weighted images were concatenated across three scans, rotated to standard orientation (FSL, reorient2std), motion- and distortion-corrected (FSL) as in (144). We then computed the diffusion tensor invariants (e.g., fractional anisotropy, FA, mean diffusivity, MD) and $S_0$ image (employed as $T_2$-weighted MRI) using FSL (dtifit).

After reorienting the $S_0$ image to a standard orientation (FSL, reorient2std), and performing bias field correction (SPM8, London, UK), we computed the affine transformation that aligned it to the $T_1$wEPI image. For each subject, the FA map, $S_0$ image and PAG delineation (see below) were then coregistered to MNI space by concatenating the affine transformation mapping the $S_0$ image to the $T_1$wEPI, with the affine and the non-linear transformations (described above) mapping the $T_1$wEPI to the 1mm-isotropic resolution $T_1$-weighted MNI152.

*Delineation of the periaqueductal gray and its subregions.* In each subject, an expert rater (M.B.) manually delineated the label (i.e., binary mask) of a cylindrical region surrounding the cerebral aqueduct (defined from the mean diffusivity image, as MD > 0.0015) and hyperintense in the $S_0$ image ($T_2$-weighted MRI), which comprised both the periaqueductal gray (PAG) and the upper part of the dorsal raphe (DR). Then based on the FA contrast, the upper DR was manually discriminated from the PAG (the DR was darker than the ventrolateral part of the PAG), and two labels (PAG, upper DR) were generated for each subject. Then, within the whole PAG label, M.B. manually delineated four bilateral PAG subregions (dorsomedial, DMPAG, dorsolateral, DLPAG, lateral, LPAG, and ventrolateral, VLPAG) in native space using a postmortem brainstem atlas (145) as a guide to define the neighborhood relationships and location of PAG subregions within the PAG. The DMPAG was defined as the column hypointense in FA posterior to the aqueduct; the DLPAG was defined as a roughly triangular prism hypointense in FA, anterolateral to DMPAG and extending only in the upper part of the PAG; LPAG was identified as a roughly triangular prism hypointense in FA, lateral to the aqueduct and anterolateral to DLPAG; finally, VLPAG was defined as a roughly triangular prism slightly hyperintense compared to the other subregions, anterolateral to LPAG and extending only in the lower part of the PAG. Single-subject labels of the whole PAG and PAG subregions were coregistered to MNI space as explained above. Then, we computed the spatial overlap (range: 0-100%) of these labels across subjects to yield probabilistic atlas labels of the PAG and its subregions in MNI space. Each probabilistic atlas label was validated by computing its *internal consistency* across subjects, as the modified Hausdorff distance between each label and the probabilistic atlas label (thresholded at 35%) generated by averaging the labels across the other subjects (leave-one-out cross validation). Note that due to the limited coverage of the diffusion weighted imaging protocol only the upper part of the DR was covered in this study; thus, DR delineations were not used to generate a DR probabilistic atlas label. Rather, M.B. manually improved the semi-automatic delineations of DR of (144) to ensure contiguity between them and the newly generated PAG probabilistic label, which yielded an updated DR probabilistic label.

*Functional connectivity analysis.* To avoid Type II error from stringent group-level threshold that decrease the ability of detecting small, but reliable connectivity signals (146), we opted to separate signal from noise using a bootstrapping analysis. We randomly resampled 80% of the sample ($N = 72$) 1000 times. In each iteration, we estimated cortical connectivity using a combination of surface- and volume-based analyses, as outlined in (Kleckner et al., 2017). Specifically, on the subject level, we ran a voxel-wise regression on left and right hemispheres of MNI152 and subcortical volume of MNI305 to compute the contrast effect size (contrast * beta) of the seed time series. On the group level, we concatenated the contrast effect size maps from all subjects and ran a general linear model analysis to test whether the group mean differed from zero (two-tailed one-sample t-test). This yielded final group maps that showed regions whose fluctuations significantly correlated with the seed's BOLD time series. We binarized the group maps to retain positive connectivity surviving the liberal threshold of $p < .05$ and summed the binarized maps across all 1000 iterations to obtain a 'bootstrapped' connectivity map that ranges between 0-1000.

*Evaluating the optimal number of clusters (k) for k-means clustering analyses.* We ran k-means clustering analyses with a range of *k values* ($k = 2$ to 6 for seven cortical seeds and $k = 2$ to 13 for 14 subcortical seeds). For both sets of analyses, we used 10 initializations of new centroid positions with a maximum of 1000 iterations each to find the lowest local minimum for sum of distances (*kmeans,* MATLAB). We calculated the Calinski-Harabasz Criterion (121) for each analysis (*evalclusters*, MATLAB). A higher Calinski-Harabasz Criterion indicates larger between-cluster variance and smaller within-cluster variance (i.e., a better clustering solution). Cortical maps derived from cortical seeds yielded in an unambiguously optimal solution with $k = 2$ based on the Calinski-Harabasz Criterion (**Supplementary Figure 9A**). For cortical maps derived from subcortical seeds, the Calinski-Harabasz Criterion remained stable for solutions between $k = 2$ to 9 but showed a drastic increase at $k > 10$ (**Supplementary Figure 9B**). To optimize parsimony (i.e., preferring a solution with fewer clusters when the evaluation criterion is similar) and consistency between analyses (i.e., aligning with the previous two-cluster solution for cortically seeded maps), we investigated the conjunction maps for $k = 2$ to 4 for interpretability.

*Comparing k-means solutions for interpretability.* The two-cluster solution differentiated the discovery maps with sparse cortical connectivity from those showing more widespread cortical connectivity (rather than differentiating spatially distinct connectivity patterns per se). The cluster with widespread connectivity included discovery maps from larger seeds in the mdThal, LGN, hippocampus, dAmy, NAcc, SC, SN and VTA and showed connectivity to large swaths of the cingulate (subgenual, pregenual, anterior mid, and posterior), insular (anterior to posterior), and frontal (medial prefrontal and superior frontal) cortices. It was observed in the three-cluster solution as well (Cluster 3; **Supplementary Figure 4**). The sparse cluster from the two-cluster solution was further divided into two almost non-overlapping clusters in the three-cluster solution, one that included maps from seeds in the lower brainstem (LC, PBN, VSM), which primarily showed connectivity to the posterior cingulate cortex, supramarginal gyrus and some medial and lateral occipital regions (Cluster 1; **Supplementary Figure 4**), and one that included maps from seeds in the upper brainstem (PAG, DR) and the hypothalamus, which showed connectivity to the aMCC and parahippocampal gyrus (Cluster 2; **Supplementary Figure 4**). The four-cluster solution yielded the same clusters as the three-cluster solution, with the exception that the hippocampal seeded map split off from the dense cluster to form its own cluster and did not further improve solution interpretability.

*Ruling out partial volume effects.* Given the size of the brainstem seeds in relation to the voxel size, we sought to rule out partial volume effects. First, we eroded each brainstem seed by one voxel. For five seeds, eroding by one voxel reduced the seed to less than ten total voxels: VTA, DR, PBN, LC, and VSM. For these seeds, we dilated neighboring seeds by one voxel and subtracted those voxels from the target seed. Thus, an eroded VTA seed was created by subtracting dilated hypothalamus and SN seeds, an eroded DR seed was created by subtracting a dilated PAG seed, an eroded PBN seed was created by subtracting a dilated LC seed, and an eroded LC seed was created by subtracting a dilated PBN seed. VSM neighbored no other seeds. The time series of the eroded seeds were highly correlated with the time series of the full seeds (*M rs* = .74 - .99; *s.d. rs* = .007 - .11), suggesting that the signals from the outer most voxels of the seeds were not significantly impacted by potential noise introduced by partial volume effects. For this reason, we ruled out the possibility that partial volume effects contaminated functional connectivity between the brainstem seeds and the rest of the brain.
